# Establishment, conservation, and innovation of dorsal determination mechanisms during the evolution of vertebrate paired appendages

**DOI:** 10.1101/2025.07.02.662459

**Authors:** M. Brent Hawkins, Sofía Zdral, Silvia Naranjo, Miguel Juliá, Manuel Sánchez-Martín, Jacob M. Daane, Nicolas Cumplido, David Jandzik, Daniel M. Medeiros, Sarah K. McMenamin, Matthew P. Harris, Juan J. Tena, Marian A. Ros

## Abstract

Limb function requires polarized anatomy across the dorsal-ventral (DV) axis, but it is unclear when the capacity for DV differentiation of paired appendages arose in evolution. Here we define ancestral DV patterning programs in the fins of fishes. We show that the orthologue of the limb dorsal determinant, Lmx1b, is required to establish dorsality in zebrafish pectoral fins and is regulated by a conserved *LARM* cis-regulatory hub. However, *lmx1bb* expression in median fins is unaffected by removal of the *LARM*, suggesting its regulation is an evolutionary innovation specific to the paired appendages. Although we find the *LARM* is highly conserved across gnathostomes, we identify specific alteration of this region in hillstream loaches, fishes which naturally parallel “double-ventral” fin phenotypes observed in *lmx1bb* and *LARM* mutants. Altogether our findings indicate *LARM*-mediated dorsal identity is an ancestral feature of paired appendages that provide a prepattern for limb evolution and lineage diversification.

## INTRODUCTION

The evolutionary transition from fins into limbs involved integrated anatomical transformations along the proximal-distal (PD), anterior-posterior (AP), and dorsal-ventral (DV) axes. While differences between fin and limb skeletons are dramatic on the PD (shoulder to finger) and AP (thumb to pinky) axes, the morphological polarity along the DV (back of the hand to palm) axis remain to be elucidated in fins, despite being critical for limb function (Haro et al., 2021). The correct arrangement of internal elements, such as flexor and extensor muscles, and superficial structures, such as hair, nails, pads, and glands, on the dorsal and ventral aspects of the limb is required to enable movement, articulation, and sensory functions (**Fig. 1a**).

**Fig. 1.**
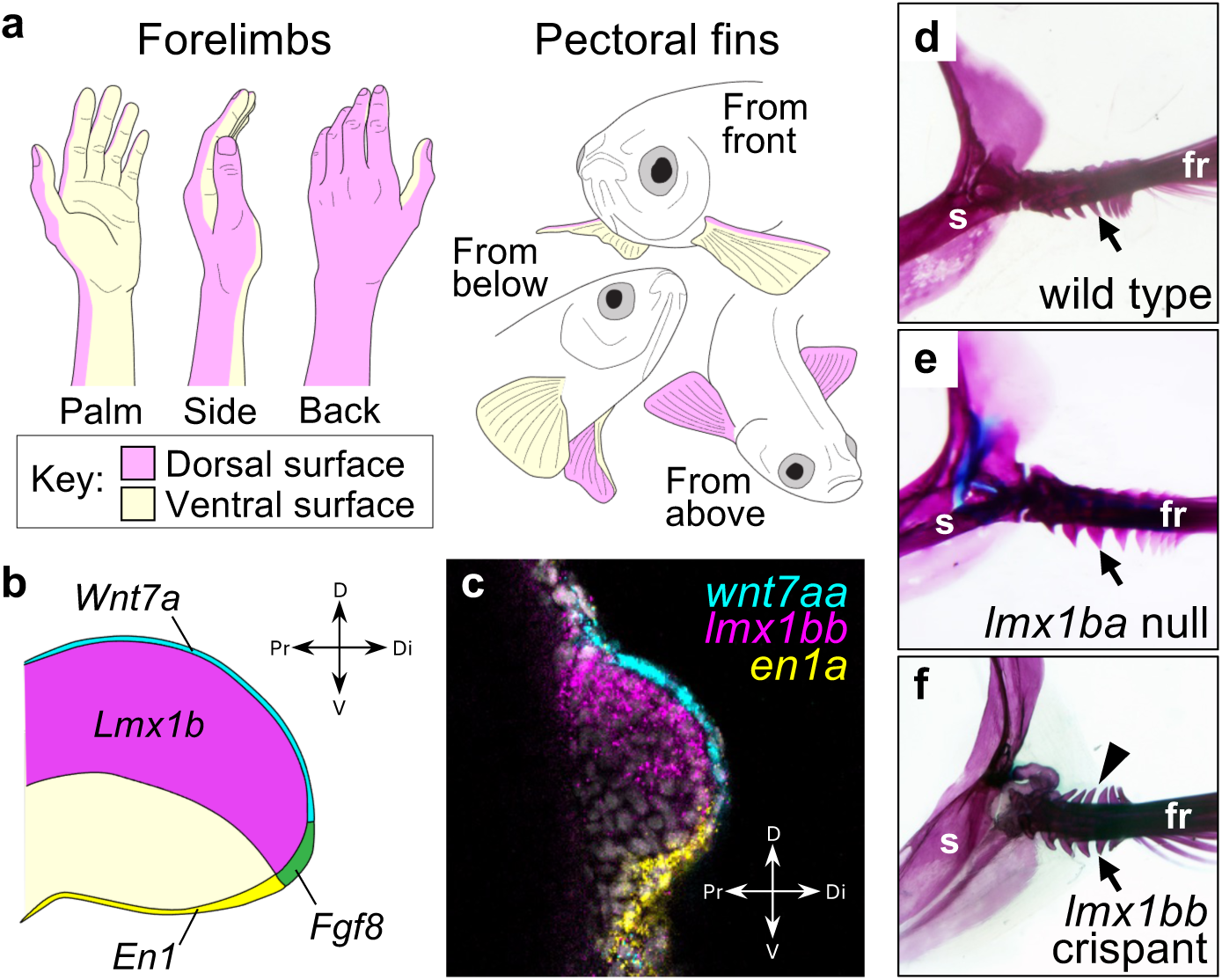
Genetic regulation of dorsal specification is functionally conserved in limb and paired fin development. **a**, Schematic of vertebrate paired appendages indicating the dorsal (pink) and ventral (yellow) regions of limbs (left) and fins (right). **b**, Diagram of DV patterning gene expression domains in the developing limb bud. *Wnt7a* (blue) in the dorsal epithelium induces the dorsal determinant *Lmx1b* (pink) in the dorsal mesenchyme while *En1* (yellow) is expressed in the ventral ectoderm. **c**, Confocal section of an HCR-labelled 36 hpf zebrafish pectoral fin bud demonstrating conserved expression patterns of DV patterning gene orthologues. **d**, Wild type adult zebrafish pectoral fin skeleton stained with alizarin red (for bone) and alician blue (for cartilage) exhibiting fin ray flanges restricted to the ventral aspect (black arrow, n=10). **e**, Null *lmx1ba* mutants have wild-type fin anatomy (n=10). **f**, The fins of *lmx1bb* crispant zebrafish express a double-ventral phenotype and develop ectopic fin ray flanges on the dorsal aspect (black arrowhead). In 45 adult *lmx1bb* crispant fish, fin ventralization was unilateral in 13 individuals and bilateral in 15 individuals. Ventralization is never observed in wild type or *lmx1ba* mutant fish. Dorsal to top, distal to right in b-f. fr, fin rays; s, shoulder.

The existence of DV polarized anatomy is less apparent in fins compared to limbs, and it is unknown whether and how DV polarity is developmentally patterned in fins. It is possible such patterning mechanisms are shared between fins and limbs due to their derivation from a common antecedent appendage, or alternatively extensive DV polarity could be a limb-specific feature. A recent indication of morphological DV asymmetry in the fin rays of tetrapomorph fish fossils, windows into the early skeletal transitions of limb-harboring vertebrates, suggests DV patterning mechanisms existed prior to the emergence of limbs (Stewart et al., 2020). However, it remains unclear if such cues are shared with ray-finned fishes and other jawed vertebrates.

The identification of the molecular factors that establish DV polarity in the developing limb was achieved through seminal experimental studies in mouse and chicken (**Fig. 1b**) (Parr and McMahon, 1995; Riddle et al., 1995; Yang and Niswander, 1995; Loomis et al., 1996; Vogel et al., 1995). Of the known genetic regulators, Lmx1b is both necessary and sufficient for the formation of limb structures with dorsal identity, while ventral identity is considered a default state (Parr and McMahon, 1995; Riddle et al., 1995; Vogel et al., 1995). Loss of *Lmx1b* function in the mouse results in an anatomically mirrored “double-ventral” phenotype in the distal limb, wherein ventral features develop on the dorsal aspect, while dorsal structures such as hair and nails are reduced or absent (Chen, Lun et al., 1998; Chen, Ovchinnikov et al., 1998). Reciprocally, loss of En1 activity in the ventral ectoderm causes a “double-dorsal” phenotype at distal levels, resulting in dorsal structures developing on the ventral side of the limb (Logan et al., 1997).

Recent studies have identified the gene regulatory architecture that controls *Lmx1b* transcription specifically in the developing mouse limb. After initial activation by Wnt7a and other signals from the dorsal ectoderm, *Lmx1b* in the dorsal mesenchyme maintains its own expression through the binding of two upstream autoregulatory elements (**Fig. 2**), called *<u>L</u>mx1b*-<u>a</u>ssociated *cis*-regulatory <u>m</u>odules 1 and 2 (*LARM1* and *LARM2*) (Haro et al., 2017; Haro et al., 2021). Based on reporter assays in the chicken embryo, *LARM1* can be divided into two elements: a putative silencer sequence (*LARM1s*) and an enhancer region (*LARM1e*). Targeted removal of all *LARM* elements (Δ*LARM1/2*, 7.6 kb) in mice recapitulates the double-ventral phenotype of *Lmx1b* loss-of-function only in the limb, therefore retaining mutant viability and demonstrating the limb-specific regulation of the *LARM* (Haro et al., 2021). The limbs of these mutants are incapable of lifting the body due to ventralization of dorsal muscular elements and severe joint malformations, demonstrating that proper DV polarity is essential for limb-based locomotion. While initial comparative bioinformatic analysis found that the *LARM* is highly conserved across tetrapods and lobe-finned fishes, orthologues in other branches of vertebrate phylogeny were not readily identified by conventional sequence conservation analyses (Haro et al., 2021).

**Fig. 2.**
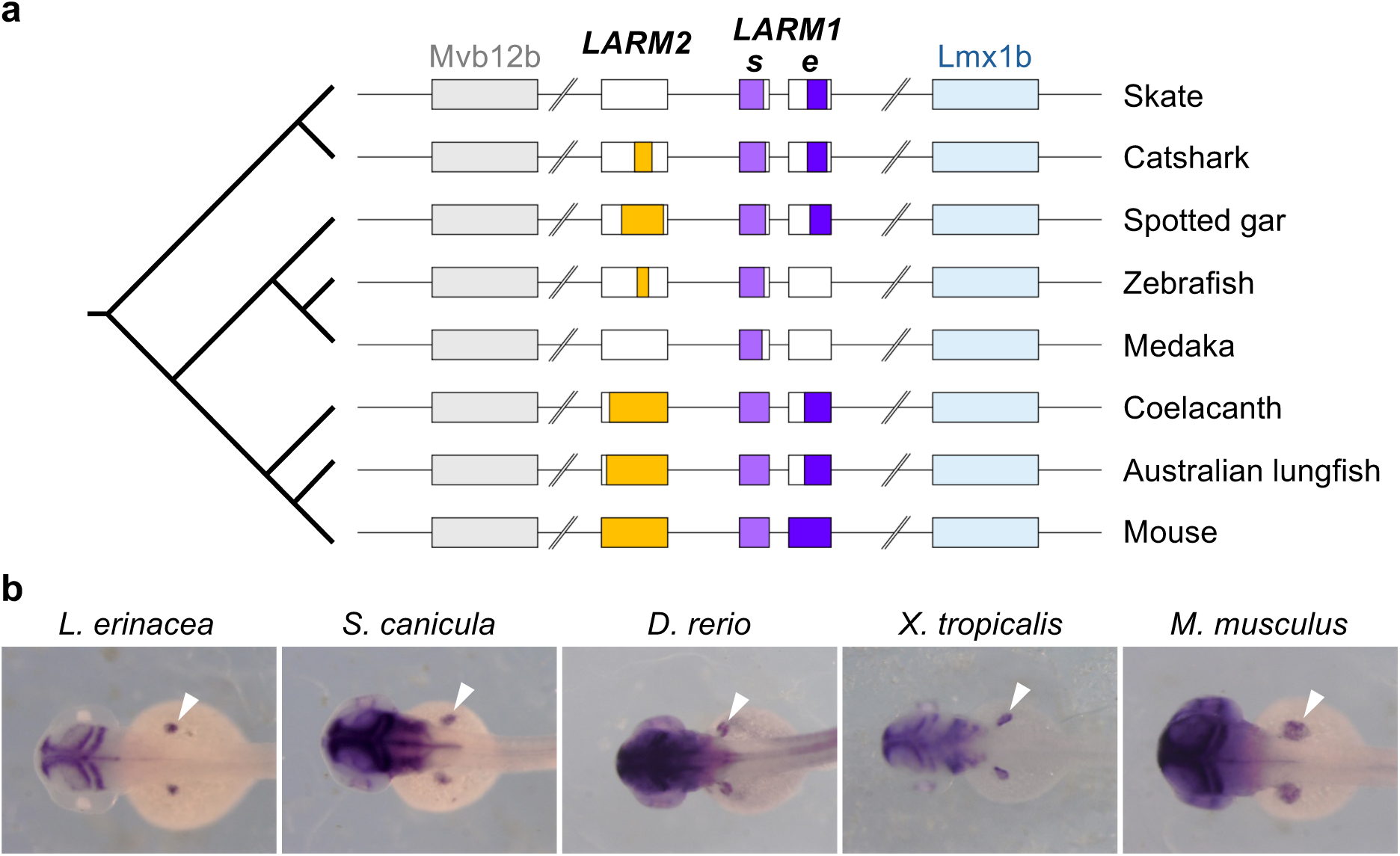
The regulatory hub that controls dorsal limb *Lmx1b* expression is conserved across jawed vertebrates. **a**, Conservation of *LARM* elements across gnathostome phylogeny. Width of colored region in each box represents extent of sequence conservation. The *bb* paralogue is shown for medaka and zebrafish. **b**, *in situ* hybridization of GFP transcripts in stable F_1_ reporter transgenics show *LARM1* elements from various gnathostome species drive expression in the zebrafish pectoral fin in transgenic assays.

While the presence of DV polarized morphology and its genetic regulation have been well described in the amniote limb, it is unclear if, and how, such polarity is established and modified in the development of fins. Did DV patterning ques arise uniquely in the evolution of the limb, or was this mechanism already present in ancestral vertebrate paired appendages and thus shared between fins and limbs? Here we describe anatomical DV polarity of paired fins in the zebrafish (*Danio rerio*) and experimentally assess genetic regulation of dorsal and ventral fate, comparing activity of teleost specific orthologues to developmental DV specification as known in the chick and mouse. We reveal broad conservation of DV patterning mechanisms in developing paired appendages as well as identify an ancient gnathostome-specific origin and shared functionality of the *LARM* elements in paired appendages. On this background of evolutionary conservation, we highlight association of natural variation in *LARM* domains within certain teleost lineages harboring pectoral fin elaborations that mirror double-ventral phenotypes. This lineage-specific variation in loaches is the first indication of adaptive DV alteration in skeletal patterning and provides a clear genomic signature of its alteration in the *LARM* regulatory hub.

## RESULTS

### A dorsal determinant program shared across fins and limbs

Mouse and chick limb buds express DV patterning factors in conserved domains during their development (**Fig. 1b**), with *Wnt7a* in the dorsal epithelium subtended by *Lmx1b* in the dorsal mesenchyme, and *En1* in the ventral ectoderm (Dealy et al., 1993; Parr and McMahon, 1995; Riddle et al., 1995; Yang and Niswander, 1995; Loomis et al., 1995; Vogel et al., 1995). To assess similarities in temporal and spatial expression of fish orthologues, we performed RNA-fluorescence in situ hybridization with amplification by hybridization chain reaction (HCR™ RNA-FISH) in the developing fin buds of zebrafish embryos. Our analysis reveals that at 36 hours post-fertilization (hpf), a stage comparable to early budding limb buds, orthologues of these genes, *wnt7aa*, *lmx1bb*, *and en1a*, are expressed in these same conserved domains as seen in mouse and chick (**Fig. 1c**). These results agree with reports of polarized expression based on previous *in situ* hybridizations in zebrafish (Ekker et al., 1992; Reifers et al., 1998; Neumann et al., 1999; Grandel et al., 2000; Uemura et al., 2005). Because *lmx1bb* has been reported in the adductor muscle of the zebrafish pectoral fin bud (Uemura et al., 2005), whereas in mouse *Lmx1b* is restricted to lateral plate mesoderm derivatives and excluded from the muscle lineage (Schweizer et al., 2004), we asked whether the expression domains of *Lmx1b* orthologues differ between species. HCR™ RNA-FISH using probes for *lmx1bb* and two muscle lineage markers, *lbx1b* (Jagla et al., 1995) and *pax3a* (Seger et al., 2011; Minchin et al., 2013), revealed non-overlapping expression in fins, supporting conserved mesenchymal-specific expression of *lmx1bb* in fins as in limbs (**Supplementary Fig. 1**).

### Functional integration of dorsal fate regulation in zebrafish

To determine if these genes share a conserved function in the DV patterning of the fin, thus predating the fin-to-limb transition, we used CRISPR-Cas9 to generate loss-of-function alleles of the zebrafish orthologues of *En1* (*en1a*, *en1b*) and *Lmx1b* (*lmx1ba*, *lmx1bb*) (**Supplementary Table 1**) and analyzed their effect on the fin phenotype. Wild type zebrafish have a DV asymmetry, developing bony flanges only on the ventral side of the pectoral fin (**Fig. 1d**). Consistent with the observation that *lmx1ba* is not detected in the zebrafish pectoral fin buds (Burzynski et al., 2013), homozygous mutants for *lmx1ba* are phenotypically wild type and show no modifications to DV polarity (**Fig. 1e**). However, crispant zebrafish injected with a CRISPR-Cas9 guide targeting the *lmx1bb* coding region show a dramatic “double-ventral” fin phenotype in which the dorsal aspect of the pectoral fin is transformed to develop ventral anatomy including the bony flanges (**Fig. 1f**). This phenotype parallels the double-ventral limbs of *Lmx1b* null mice (Chen, Lun et al., 1998). Similar to the neonatal lethality observed in *Lmx1b* null mutants, homozygous loss-of-function of *lmx1bb* is incompatible with viability past larval stages in the zebrafish (Schibler and Malicki, 2007), preventing assessment of the adult fin phenotype. Our results demonstrate that, as in limbs, Lmx1b is required to establish dorsal specification. This function is retained in the *lmx1bb* paralogue, as *lmx1ba* loss-of-function does not have discernable fin phenotype nor is it able to complement the loss of *lmx1bb*.

As En1 has been shown to dorsally restrict *Wnt7a* and *Lmx1b* expression in amniote limbs, we asked if this genetic epistasis at the core of DV axis patterning is also observed during fin development. Zebrafish *en1a*;*en1b* double-null animals are adult viable and have a pigment patterning phenotype confirming loss of En1 function (**Supplementary Fig. 2**). In contrast to the *En1* knockout mouse, we do not observe a dorsalization phenotype, such as loss of ventral flanges, in the proximal dermal rays and these rays retain wild-type flange morphology. As the ventral surface does not have obvious polarized structures apart from the proximal flanges, changes in more distal fates in the fin are not able to be scored. In mouse, loss of *En1* function results in the activation of *Wnt7a* expression in the ventral ectoderm (Loomis et al., 1996), which in turn induces ectopic *Lmx1b* expression in the ventral mesenchyme. At 36 hpf we do not observe significant alteration in the extent of *wnt7aa* or *lmx1bb* expression in *en1a*;*en1b* double mutant fin buds (**Supplementary Fig. 2f).** However, by 56 hpf mutant fins exhibit ectopic ventral expression of *wnt7aa* and *lmx1bb* transcripts (**Supplementary Fig. 2h**). Altogether our results demonstrate the integrated signaling network of DV specification as defined in mouse limb is intact and functionally conserved in zebrafish pectoral fins.

### Ancestral origin and evolutionary conservation of *Lmx1b* regulatory elements

Given the conserved epistatic interactions between DV patterning molecules in fins and limbs, we asked if transcriptional control of *lmx1bb* in the fin is controlled by similar regulatory logic through *LARM* orthologues. Previous analysis found that the *LARM* elements are conserved among tetrapods as well as the coelacanth, but orthologs of this regulatory hub could not be found in non-sarcopterygian fishes by sequence identity (Haro et al., 2021). In the mouse genome, the *LARM* is situated in a ∼6 kb interval located ∼60 kb upstream of the *Lmx1b* transcription start site*. Lmx1b* is the only coding gene within a Topologically Associated Domain (TAD) that spans from the 3’ end of the upstream *Mvb12b* gene to the promoter of the downstream *Zbtb43* gene (Supplementary Fig. 3a). We find that this syntenic relationship is maintained across gnathostomes and, to some extent, also preserved in cyclostomes which lack paired fins (**Supplementary Fig. 3b**).

To identify putative *LARM* orthologs in species outside of the lobe-finned fish lineage, we generated a Hidden Markov Model (HMM) using the multiple alignment of *LARM* sequences from 12 vertebrate species (chicken, turkey, coelacanth, human, bat, whale, cow, manatee, otter, seal, gecko, and mouse). We then used this HMM to search for *LARM* elements in the genomes of fish species representing major vertebrate lineages as well as key phylogenetic positions that bracket the fin-to-limb transition. Using this approach, we identified orthologous *LARM1* and *LARM2* elements across jawed vertebrate lineages with varying degrees of sequence conservation (**Fig. 2a**). However, analysis of *Lmx1b* loci in cyclostomes, the lamprey and hagfish, found no *LARM*-like regions. Among gnathostomes, the *LARM1s* element exhibits the highest sequence conservation across groups and was identified in each species examined suggesting that it represents a core functional component of the locus. In cartilaginous fishes, the *LARM1s* and *LARM1e* elements were found in both skate and catshark, but a partial *LARM2* element could be identified only in the catshark. Consistent with previous findings and a more proximate evolutionary distance, *LARM1e* and *LARM2* elements were also highly conserved in the non-tetrapod sarcopterygians the coelacanth and the Australian lungfish. Among ray-finned fishes, *LARM1e*, *LARM1s*, and *LARM2* orthologs were identified in the sterlet and spotted gar but teleosts exhibited reduced conservation of these elements. While *LARM1s* and a partial (non-significant) *LARM2* sequence were identified in zebrafish, medaka only retains the *LARM1s* element, and neither species had a recognizable *LARM1e* by sequence identity according to our HMM. Interestingly, in both zebrafish and medaka, these elements are only found upstream of the *lmx1bb* paralogue and not *lmx1ba*, indicating that paralogue sub-functionalization of DV patterning activity occurred following the teleost-specific whole genome duplication. This is consistent with our finding that loss of *lmx1ba* had no effect on fin DV patterning (**Fig. 1e**). Altogether these results demonstrate that the *LARM1e*, *LARM1s*, and *LARM2* elements were all present in the common ancestor of jawed vertebrates and have subsequently diverged at the sequence level across gnathostome phylogeny.

To determine if non-sarcopterygian *LARM* elements can drive expression in the paired appendages, we used reporter transgene assays in zebrafish using constructs wherein different *LARM* orthologues (**Supplementary Table 2**) were cloned upstream of a minimal promoter driving GFP expression. Using this approach, we found that all *LARM* orthologues tested were sufficient to drive reporter expression in a similar pattern in the developing pectoral fins in transgenic zebrafish, at both the transcript (**Fig. 2b**) and protein levels (**Supplementary Fig. 4**), demonstrating their functional conservation across gnathostome lineages. Additionally, these results indicate that the *LARM1s* ortholog, a putative silencer in mouse based on chicken enhancer reporter assays (Haro et al., 2021), activates transgene expression in zebrafish (**Fig. 2b**). This suggests that local genomic context in which the *LARM* acts may impact their effect on transcription.

### *LARM* function is conserved in the development of fins and limbs

In mouse, the *LARM* elements comprise a limb-specific regulatory hub of *Lmx1b*, and targeted homozygous removal of *LARM1* and *LARM2* recapitulates the double-ventral limb phenotype observed in *Lmx1b* null mutants while retaining viability (Haro et al., 2021). We asked if the zebrafish element identified by our HMM search, a *LARM1s* orthologue of the *lmx1bb* locus 164 bp in length, shares similar regulatory function. We used CRISPR-Cas9 to generate stable mutant lines in which this element was removed from the zebrafish genome. Surprisingly, homozygous *LARM1s* deletion mutants were phenotypically wildtype and exhibited no defects in pectoral fin DV patterning, suggesting that this element alone is not necessary for paired appendage expression of *lmx1bb* (**Supplementary Fig. 5**). Concomitantly, we found that this 164 bp zebrafish element was unable to rescue normal DV patterning when it replaced the endogenous *LARM1* and *LARM2* elements in transgenic mice (**Fig. 3a**). In this experiment, we substituted the endogenous 7.6 kb region encompassing *LARM1* and *LARM2* in the mouse genome with the 164 bp zebrafish *LARM1s* conserved element, which neither rescued the double ventral phenotype of forelimbs or hindlimbs, nor *Lmx1b* expression (**Fig. 3b,c**). A consistent partial rescue was observed at the elbow, which no longer exhibited the typical dislocation seen in Δ*LARM1/2* homozygous mutants lacking all *Lmx1b* expression in the limbs (**Supplementary Fig. 5**). Strikingly, supplementing the endogenous mouse *LARM* with two copies of the 164 bp zebrafish *LARM1s* zebrafish element produced a distal double-dorsal phenotype even in heterozygosity (**Fig. 3d,e**). The extra regulatory input from the tandem zebrafish elements caused ectopic *Lmx1b* expression in the distal ventral mesoderm (**Fig. 3f**), however this occurred without alterations in the normal expression patterns of *En1* or *Wnt7a*. Thus, even though the zebrafish *LARM1s* minimal element does not fully functionally complement the endogenous mouse *LARM*, the element may act as an enhancer in varied genomic contexts and with dose-dependent effects. Interestingly, transgenesis assays show that the zebrafish *LARM1s* element contains regulatory information to drive expression in the pectoral fin (**Fig. 2b**). When two copies of this sequence were injected in tandem in transgenesis assays, fin GFP signal was increased over the single copy construct (**Supplementary Fig. 4**). This suggests a possible booster function of the zebrafish 164 bp *LARM1s* element, increasing the activity of other nearby enhancers, and provides a potential explanation of the ventrally-expanded *Lmx1b* expression and double-dorsal phenotype obtained when two copies of this sequence were introduced in the mouse *Lmx1b* locus (**Fig. 3e,f**).

**Fig. 3.**
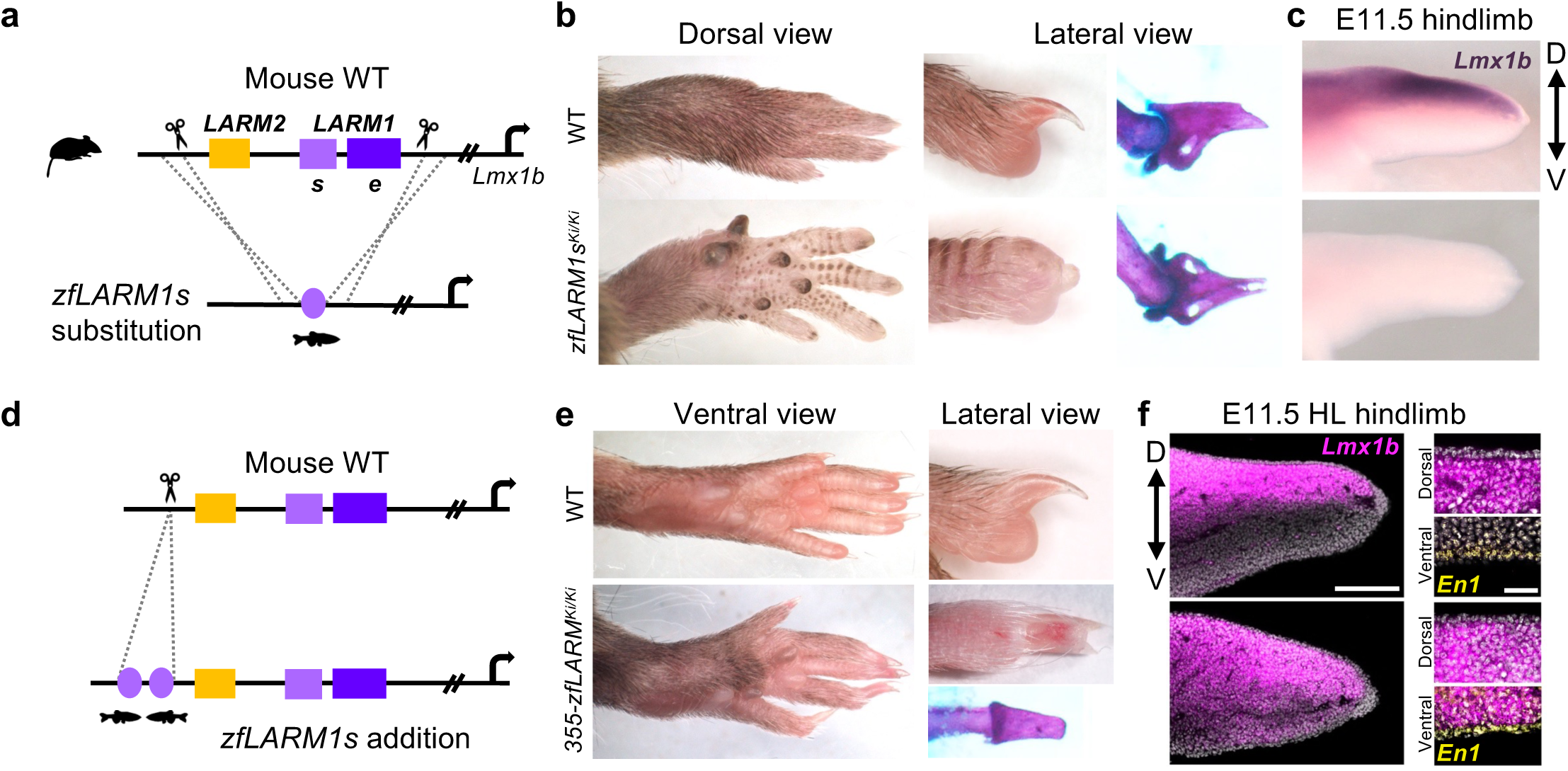
The zebrafish *LARM1s* element is not sufficient to drive dorsal fates in mice but has additive effects with mouse *LARM*. **a**, Schematic of transgenic design to test the activity of zebrafish *LARM1s* conserved element (164 bp) in place of the endogenous *LARM* region (7.6 kb) in mouse limb patterning. **b**, The zebrafish *LARM1s* element is not sufficient to drive *Lmx1b* expression and the development of dorsal features as *zfLARM1s^Ki/Ki^*mice have double-ventral limbs and **c**, loss of *Lmx1b* expression in the dorsal limb mesenchyme. **d**, Schematic of identified *355-zfLARM ^Ki^* allele in which two *zfLARM1s* elements are incorporated in addition to the endogenous mouse *LARM1* and *LARM2*. **e**, The limbs of *355-zfLARM ^Ki/Ki^*mice express a double-dorsal phenotype and **f**, display ectopic ventral expression of *Lmx1b* while *En1* is maintained in the ventral ectoderm.

To extend our analysis into the conservation of *Lmx1b* regulation, we considered the contribution of the partial *LARM2* element identified in zebrafish by our HHM search. Inspection of the *lmx1bb* regulatory landscape in zebrafish ATACseq data sets from 48 hpf embryos revealed several proximal open chromatin peaks. The most prominent peak corresponds to the *LARM1s* ortholog, while another smaller peak overlaps with the partially conserved *LARM2* element, suggesting a larger regulatory region than that defined by sequence identity alone **(Fig. 4a)**. Additionally, VISTA genomic alignments of the *Mvb12b*-*Lmx1b* intergenic region using the “Holostean bridge” approach (Braasch et al., 2016) revealed distal sequences with low sequence similarity to the *LARM2* region (**Supplementary Fig. 6**). Considering this, we overlayed the full ∼6 kb mouse *LARM* region onto the zebrafish locus using the 164 bp element as an anchoring point and selected a ∼6 kb region that included the putative *LARM2* region. Stunningly, homozygous deletion of this 6.8 kb holo-*LARM* interval recapitulated the double-ventral fin phenotype observed in crispant animals while avoiding the lethality observed in *lmx1bb* null zebrafish (**Fig. 4b**). These results indicate that the zebrafish *LARM* hub retains a conserved regulatory function despite extensive sequence divergence from sarcopterygian orthologues.

**Fig. 4.**
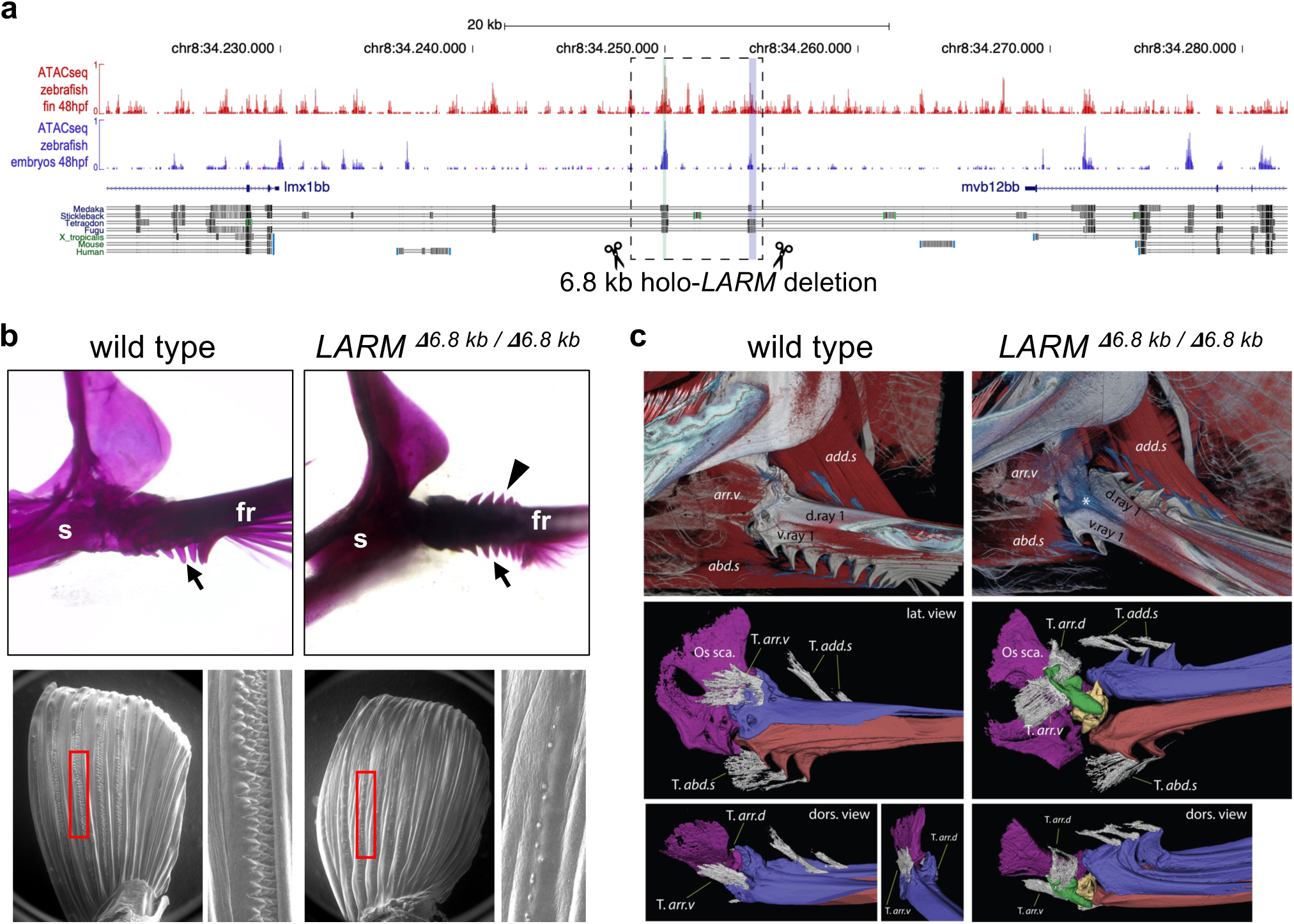
*LARM*-regulated *Lmx1b* directed specification of dorsal fate is ancestral feature of paired appendages. **a**, ATAC-seq profiling reveals open chromatin at the zebrafish *LARM1s* (green bar) and cryptic *LARM2* (blue bar) orthologues. The 6.8kb holo-*LARM* deletion includes both peaks (dashed box). **b**, *LARM ^6.8kbΔ/6.8kbΔ^* mutants develop double-ventral fins with ectopic dorsal flanges (black arrow, n=10). SEM micrographs reveal wild type males develop tubercle tracts on the pectoral fin dorsal surface (bottom left; n=10), but *LARM ^6.8kbΔ/6.8kbΔ^* males do not (bottom right), consistent with a dorsal to ventral transversion of the integument (n=10). **c**, Transformation of the dorsal musculoskeletal system to the ventral configuration in *LARM ^6.8kbΔ/6.8kbΔ^* mutants and the formation of a novel anterior element (green) revealed by immunolabelling (n=5). abd.s, abductor superficialis; add.s, adductor superficialis; arr.d, arrector dorsalis; arr.v, arrector ventralis; d.ray 1, dorsal hemiray 1; Os sca, scapula; v.ray 1, ventral hemiray 1; T.*, tendon of.

The stable 6.8 kb holo-*LARM* deletion line permitted more thorough anatomical characterization of the double-ventral fin phenotype beyond what was possible with the genetically mosaic *lmx1bb* crispant animals. In addition to dorsal duplication of the bony fin ray flanges, we also observed transformations of the dorsal integument and muscular system to take on ventral fates. Adult male zebrafish develop patches of keratinized breeding tubercles on the dorsal surface of the pectoral fin (Kang et al., 2013; McMillan et al., 2013). Consistent with a transversion of dorsal to ventral identity of the integument, homozygous holo-*LARM* deletion males failed to develop tubercle patches and formed only few scattered individual tubercles (**Fig. 4b**); these males are accordingly unsuccessful in mating. The fins of holo-*LARM* deletion animals exhibit the full transformation of dorsal structures into ventralized anatomy, leading to changes in the musculo-skeletal integration of the new dorsal flanges of the rays (**Fig. 4c**). The dorsal duplication of fin ray flanges was associated with an integrated change in the insertion of the dorsal tendons coming from the *adductor superficialis*—the primary muscle that elevates and retracts the fin. In wild-type animals, the tendons insert distally in the dorsal surface of the ray, but in the holo-*LARM* deletion animals the dorsal *adductor superficialis* tendons actually insert more proximally on the ectopic dorsal flanges (**Supplementary Fig. 7, Fig. 4c**). Interestingly, this change did not involve a change in the morphology or integrative capacity of the muscle, suggesting that the dorsal phenotype was restricted to tendon and skeletal changes, consistent with derivation of these latter tissues from lateral plate mesoderm separate from muscle progenitors (**Supplementary Fig. 1**).

The most prominent change in the homozygous holo-*LARM* deletion mutants is the morphology of the most anterior first fin ray. In wild type animals, the dorsal component of the first ray (dorsal hemiray) receives the tendinous insertions from two opposite sides of the fin, the *arrector dorsalis* and *arrector ventralis*, and articulates with the scapula. Together this musculoskeletal system acts as a functional unit that controls the leading edge of the fin and moves the first ray either dorsally or ventrally. The holo-*LARM* deletion mutants, however, lose this function as the dorsal hemiray does not grow ventrally over the scapula, nor does it receive the tendinous attachments of the *arrector* muscles. These muscles instead attach to a novel bone located over the anterior side of the scapula and impair proper articulation of the fin (**Fig. 4c**). Consequently, homozygous holo-*LARM* deletion mutants are unable to retract the fin to the body and are conspicuous in the tank as the swim with their fins extended constantly outward. These phenotypic changes in the skeleton, tendons, integument, and articulation match those observed in *LARM* deletion mice, and demonstrate that *LARM*-mediated control of *Lmx1b* to specify dorsal appendage fate is a universal feature of paired appendages that arose prior to the origin of limbs.

### *LARM* and the emergence of a paired appendage-specific regulatory program

As median appendages appear before paired appendages in the vertebrate fossil record, it is hypothesized that fin patterning programs were first assembled in the midline fins and subsequently co-opted to form the paired appendages later in evolution (Freitas et al., 2006; Dahn et al., 2007; Letelier et al., 2018; Letelier et al., 2021; Hawkins et al., 2022). Recent work has found that *Lmx1b* orthologues are expressed in the posterior mesenchyme of developing dorsal and anal fins of multiple gnathostome lineages, suggesting this could be an ancestral feature of vertebrate fins that predates the origin of paired appendages (Zdral et al., 2025). To determine if median fin *Lmx1b* expression is conserved in cyclostomes, a vertebrate lineage that diverged from gnathostomes prior to the evolution of paired appendages, we assessed the expression of the two *Lmx1b* orthologues in the sea lamprey *Petromyzon marinus*. Embryonic expression of these orthologues, *Lmx1bα* and *Lmx1bβ*, in the central nervous system and ear is consistent with conserved roles in gnathostome development (**Supplementary Fig. 8**). In the midline fins, we found that both paralogues are expressed in the posterior/caudal domain of the second dorsal fin of advanced ammocoete larvae (**Fig. 5a**), again conserved with gnathostome median fins.

**Fig. 5.**
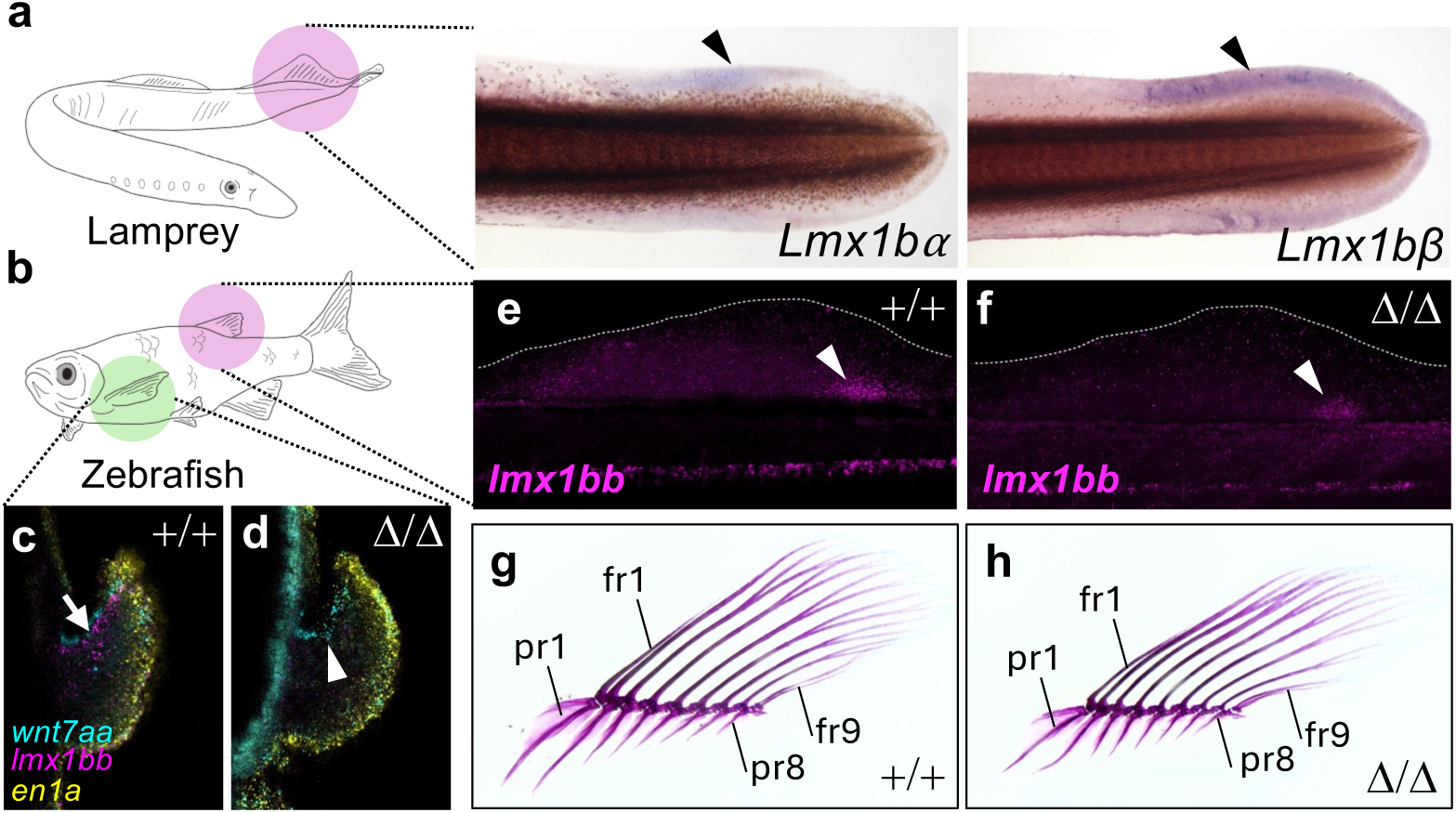
Expression of *Lmx1b* orthologues in the unpaired fins is an ancient vertebrate feature but does not require *LARM*-mediated regulation in gnathostomes. **a,** Colorometric *in situ* hybridization of *Lmx1b* orthologue expression (black arrowheads) in the posterior mesenchyme of the second dorsal fin (purple circle) of lamprey ammocete larvae. **b**, Jawed fishes such as the zebrafish possess paired fins (green circle) in addition to unpaired midline fins like the dorsal fin (purple circle). **c**, Wild-type pectoral fin express *lmx1bb* in the dorsal mesenchyme at 48 hpf (n=5). **d**, Deletion of the holo-LARM resulting the abrogation of *lmx1bb* expression in the pectoral fin (white arrowhead; n=5). **e**, In 14 dpf wild type fish, *lmx1bb* is expressed in the posterior mesenchyme of the dorsal fin bud (white arrowhead). **f**, Dorsal fin expression is maintained in *LARM ^6.8kbΔ/6.8kbΔ^* mutants. **g**, Wild-type adult dorsal fin skeleton with 8 radials and 9 rays (n=10). **h**, *LARM ^6.8kbΔ/6.8kbΔ^* mutants exhibit the wild-type dorsal fin pattern (n=10). Dorsal to left, distal to top in c,d; anterior to left, dorsal to top in a, e-h. fr, fin ray; pr, proximal radial.

To determine if *LARM-*mediated *Lmx1b* regulation is a universal feature of vertebrate appendages, or if *LARM* control is specific to the paired appendage domain, we examined *lmx1bb* expression in holo-*LARM* deletion zebrafish. Using HCR, we found that *lmx1bb* transcripts were lost in deletion mutant pectoral fins at 48 hpf (**Fig. 5c,d**). However, *lmx1bb* expression domains in other embryonic regions were unaffected (**Supplementary Fig. 9**). In the unpaired fins, we found that *lmx1bb* was normally expressed in the developing dorsal fin buds of zebrafish at 14 dpf, with a similar posterior polarity seen in lamprey and in jawed fishes (Zdral et al., 2025) rather than the dorsal polarity observed in paired fins (**Fig. 5e**). This shared expression pattern between a teleost and a cyclostome suggests that *Lmx1b* expression with a posterior polarity is an ancestral feature of vertebrate median fins. We asked if *LARM*-mediated *lmx1bb* expression also functions in median fin patterning, which would be consistent with the origin of a universal appendage enhancer function that predates the paired appendages. However, in contradiction to such a pan-fin regulation model, we discovered that dorsal fin *lmx1bb* expression was unaffected in homozygous holo-*LARM* deletion zebrafish (**Fig. 5f**), supporting the paired-appendage specific nature of *LARM* regulation. The skeletal form of the dorsal fin in the mutants was also unaffected (**Fig 5h**). These data suggest that, unlike other appendage patterning programs such as nested Hox expression (Freitas et al., 2006) and ZRS-mediated Shh patterning in the ZPA (Letelier et al., 2018), dorsal determination by *LARM*-mediated Lmx1b expression arose as a novel mechanism in paired appendages, evolving independently from median fin patterning programs specifically within gnathostomes.

### Evolutionary diversification in form and changes in DV regulation

Our data suggest broad conservation of not only signaling integration among dorsoventral regulators but also a core enhancer complex, the *LARM*, to mediate polarized expression of the key dorsal determinant *Lmx1b* during paired appendage development. It is noteworthy that, given the extensive variation in fin form across teleost diversity, as well as the importance of limb skeletal variation in shaping of the limb in tetrapod radiation, that variation in DV aspects of the limb skeleton are constrained; to date, few to no cases of lineage-specific variation in the DV polarity of paired appendages have been described or studied. In a survey of teleost pectoral fin variation, however, we identified a lineage-specific shift in DV patterning.

Hillstream loaches (families Balitoridae and Gastromyzontidae) have adapted to living in fast-flowing water and develop modified pectoral and pelvic girdles that permit them to cling tightly to rocks and climb waterfalls (Willis et al., 2019; Crawford et al., 2020). Strikingly, we discovered that members of these families, nested within the broader loach clade, develop extreme dorsal projections of the proximal fin rays reminiscent of the dorsal flanges we observe in the ventralized fins of homozygous holo-*LARM* deletion zebrafish (**Fig. 6c**). This ventralized condition is derived from an ancestral cypriniform configuration in which flanges are found ventrally, as in the zebrafish, with limited or no dorsal elaboration (**Supplementary Fig. 10**). Alignment of the *mvb12bb-lmx1bb* intergenic region across cypriniform genomes, including the hillstream loaches and close relatives, revealed conservation of the *LARM1s* and *LARM2* elements (**Supplementary Fig. 10**). However, a region adjacent to *LARM1s*, which we term *LARM1x*—itself largely conserved across the order Cypriniformes—shows reduced sequence conservation due to lineage-specific deletions in *Beaufortia* species that bear dorsal elaborations (**Fig. 6d**). We cloned the *LARM1x* element and confirmed the same deletion in *Sewellia lineolata*, a species that together with *Beaufortia* brackets the ancestral gastromyzontid node (Chen et al., 2022), suggesting this deletion is shared across this family. While teleosts lack a *LARM1e* element definable by sequence identity, the *LARM1x* region occupies an analogous genomic position immediately downstream of *LARM1s*. Analysis of *LARM1x* reveals these deletions in Gastromyzontidae disrupts predicted transcription factor binding sites otherwise conserved across cypriniform fishes (**Supplementary Table 3**). Additionally, we find that the *LARM2* element has undergone accelerated sequence evolution in the lineage leading to the hillstream loaches and the family Cobitidae (true loaches) (**Fig. 6e**). This corresponds to dorsal elaborations in true loaches: male loaches develop large dorsal pectoral fin ornaments called *lamina circularis*, which exhibit species-specific morphology (**Fig. 6f**; Havird and Page, 2010; Yashima et al., 2023). These findings suggest that variation in *LARM* elements have contributed to lineage-specific evolution in the DV axis of teleost paired appendages and may represent a target for adaptive variation.

**Fig. 6.**
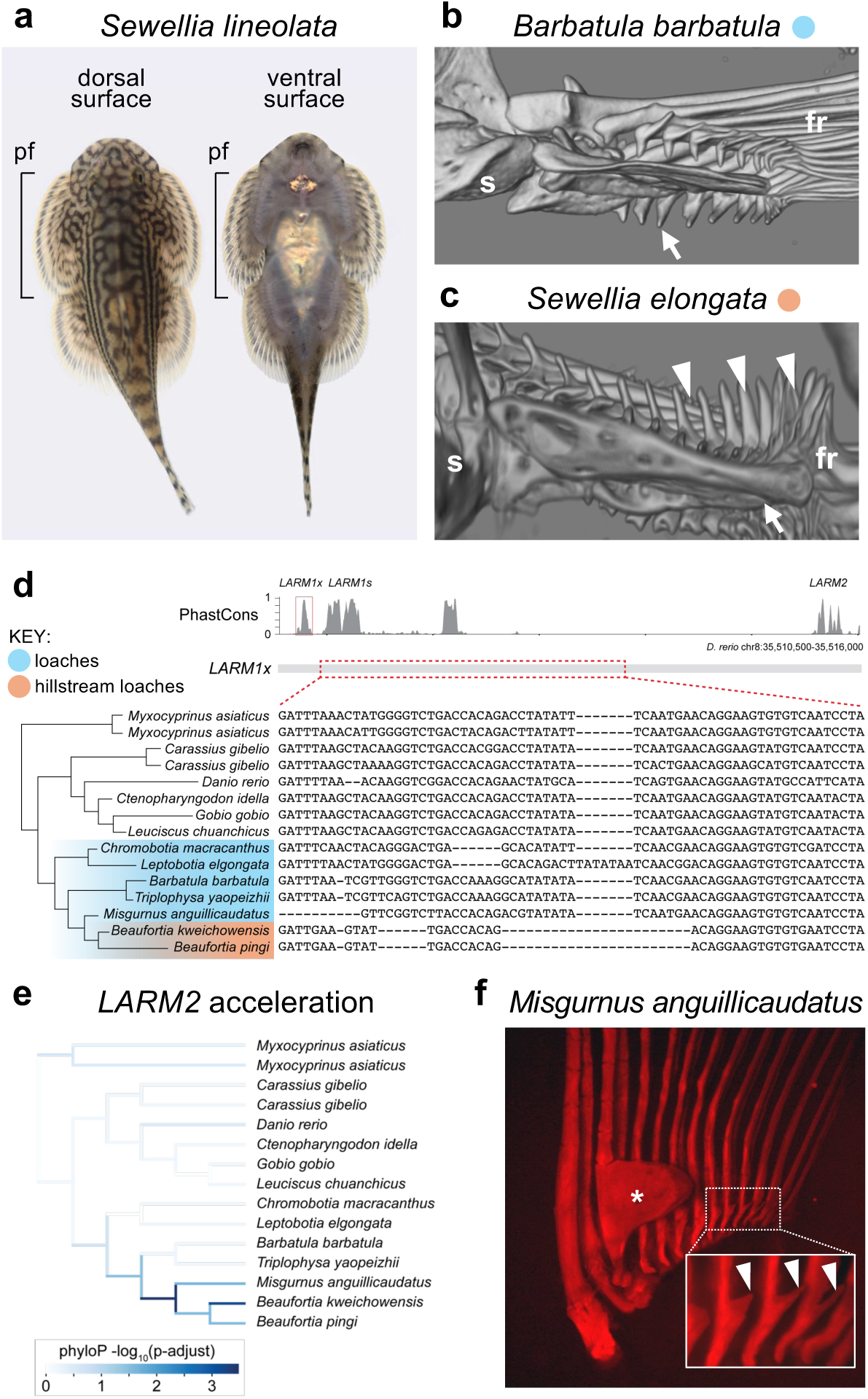
Lineage-specific evolution of the *LARM* regulatory region in the emergence of novel DV anatomies. Loaches are highlighted in blue, and hillstream loaches are highlighted in orange. **a**, *S. lineolata* demonstrating the depressed body plan and expanded fins typical of hillstream loaches. **b**, microCT reconstructed medial view of the posterior pectoral fin of *B*. *barbatula*, a loach expressing the ancestral cypriniform pectoral fin configuration with distinct ventral flanges (white arrow) and limited dorsal elaboration. **c**, Hillsteam loaches like *S*. *elongata* express a derived condition in which the dorsal rays develop large flanges (white arrowheads). **d**, Multiple sequence alignment reveals *LARM* elements are conserved across order Cypriniformes, with lineage-specific deletions in the *LARM1x* element coincident with dorsal ray elaboration. **e**, Cypriniform phylogeny mapped with sequence evolution acceleration, showing elevated rates in the *LARM2* element of cobitid and hillstream loaches. **f**, Dorsal elaborations in the pectoral fin rays of male *M*. *anguillicaudatus* include the *lamina circularis* (white asterisk) and posterior dorsal flanges (white arrowheads). Anterior to top in a; proximal to left, distal to top in b,c; anterior to left, distal to top in f. fr, fin rays; pf, pectoral fin; s, shoulder.

## DISCUSSION

Of the mechanisms that pattern the three axes of the vertebrate limb, those that determine the DV axis are the least characterized in terms of developmental genetic programs, evolutionary origins, and natural variation. Our study demonstrates that *Lmx1b* serves as the dorsal determinant in paired fins as well as limbs, and that its transcription in the dorsal mesenchyme of both appendage types is controlled by *LARM1* and *LARM2*, which comprise a conserved regulatory hub specifically devoted to the patterning of paired appendages. Results from functional manipulation of the *LARM* elements in zebrafish indicate that conservation of this dorsal program extends to at least the common ancestor of the bony fishes. The presence of *LARM* orthologues across jawed vertebrates in addition to DV-polarized *Lmx1b* expression in chondrichthyan paired appendages (Jung et al., 2018; Zdral et al., 2025) suggest that *LARM*-mediated Lmx1b dorsal specification is an ancient feature of paired fins retained from the gnathostome common ancestor. Thus, the dorsal patterning cue was in place prior to the fin-to-limb transition, and primed limbs to exaggerate DV morphological polarity beyond what is seen in fins to enable enhanced articulation and integumentary specialization.

Considering this conservation of the dorsal component of DV patterning, we were surprised to find that loss of orthologues of *En1*, which delimits *Lmx1b* activation in the limb, led to ectopic ventral *lmx1bb* expression but had no phenotypic effect on the patterning of the fin. In mouse, while Lmx1b is required for normal dorsal patterning of the zeugopod and autopod limb regions, the effect of En1 in DV patterning is limited to the autopod (Loomis et al., 1996). A plausible explanation is that zebrafish pectoral fins lack DV asymmetry at their distal extreme, preventing any possible ventral-to-dorsal transformation from being detected.

*LARM* regions are conserved across jawed vertebrates, at both the sequence and functional levels as demonstrated by our zebrafish transgenesis assays with different *LARM* orthologues. The zebrafish orthologue of the *LARM1s* element exhibited the highest conservation across lineages and was sufficient to drive transgene expression in the fin. Nonetheless, this element was unable to compensate for endogenous *LARM1* and *LARM2* in transgenic mice, and its removal from the zebrafish genome had no impact on fin DV pattern. However, removal of the 6.8 holo-*LARM* region flanking the *LARM1s* core did recapitulate the double-ventral phenotype, indicating that the zebrafish enhancer does contain this regulatory information despite sequence divergence. One interpretation of this result is that the *LARM1s* element serves as a conserved regulatory node or “tent pole” around which enhancer motifs can appear and disassemble, resulting in lineage-specific sequences and functions with reduced gnathostome-wide conservation outside of the *LARM1s* element. Concordantly, genomic alignments between mouse, gar, and zebrafish demonstrate limited conservation at large evolutionary distances (∼450 million years), while alignment of cypriniform genomes with more proximal divergence times (∼175 million years) revealed higher lineage-specific conservation.

For most developmental genetic programs involved in limb axis formation, the prevailing hypothesis is that these patterning mechanisms were first assembled in the midline fins and later co-opted in the origin of the paired fins. This framework makes the explicit prediction that paired and midline fins should exhibit similar genetic interactions between patterning molecules and shared regulatory control of gene expression. Indeed, this is the case for *Shh* in the posterior mesenchyme of both paired and midline fins, which is positively regulated by Fgf signaling and is transcriptionally controlled by the ZRS enhancer (Letelier et al., 2018; Hawkins et al., 2022). However, in the case of paired appendage *Lmx1b* regulation we found no such co-optive evolutionary link to medial fins. Rather, *LARM*-mediated *Lmx1b* transcriptional control is an innovation which arose coincident with, or subsequent to, the origin of the paired appendages, setting the stage for appendage elaboration in gnathostome lineages.

While many lineages have elaborated anatomical polarity across the dorsal and ventral appendage regions, evolutionary modification of the DV axis itself is unknown. Unlike the other spatial dimensions of limb and fin structure, variation in the DV axis has not been a source for investigation of clear adaptive variation. Our finding of a distinct phyletic phenocopy (Stebbins and Basile, 1986) in hillstream loaches of the dorsal fin ray flanges provides one such example, paralleling *lmx1bb*-deficient zebrafish. These phenotypic changes correlate with erosion of specific elements of the cypriniform *LARM* hub. These patterns highlight the potential in this locus to facilitate integrated shifts in phenotype and novel function of the fins–in this case adherence to substrates in increased flow conditions (Willis et al., 2019; Crawford et al., 2020). While it is unknown if these changes at the *LARM* hub were drivers of such change, it is clear that regulatory changes at this locus could initiate or stabilize such morphologies. As we observed in the 6.8 kb holo-*LARM* deletion fins, skeletal changes would be integrated with the musculature and facilitate novel articulation. Importantly, this shift in regulation would provide a robust developmental signal given the autoregulatory feedback observed at the *LARM*. This would provide support for evolutionary changes in the DV axis of appendage structure and facilitate natural variation within these lineages.

## ACKNOWLEDGEMENTS

The authors thank Yan Gong for assistance with SEM imaging and Stephen Treaster for curation of cypriniform genomic data.

## AUTHOR CONTRIBUTIONS

Conceptualization: MBH, SZ, MPH, JJT, MR

Methodology: All authors

Investigation: All authors

Visualization: MBH, SZ, SN, MJ, NC, DJ, JMD

Funding acquisition: DJ, DMM, MPH, SKM, JJT, MR

Project administration and supervision: MPH, JJT, MR

Writing the original draft: MBH, SZ, MPH, JJT, MR

Review writing & editing: All authors

## ETHICAL STATEMENT

All experiments involving zebrafish (*Danio rerio*) performed in this work conform to European Community standards for the use of animals in experimentation and were approved by the Ethical Committees from the Universidad Pablo de Olavide and the Andalusian Government, or Institutional Animal Care and Use Committee (IACUC) of Boston Children’s Hospital. Mouse husbandry and use were conducted according to the EU regulations and 3R principles, reviewed and approved by the Bioethics Committee of the University of Cantabria. Lamprey (*Petromyzon marinus*) care and experiments were approved by the IACUC of the University of Colorado at Boulder under protocol 2392.

## DATA AVAILABILITY

All original data are available upon request.

## CODE AVAILABILITY

All original code is available upon request.

## FIGURE LEGENDS

**Supplementary Fig. 1.**
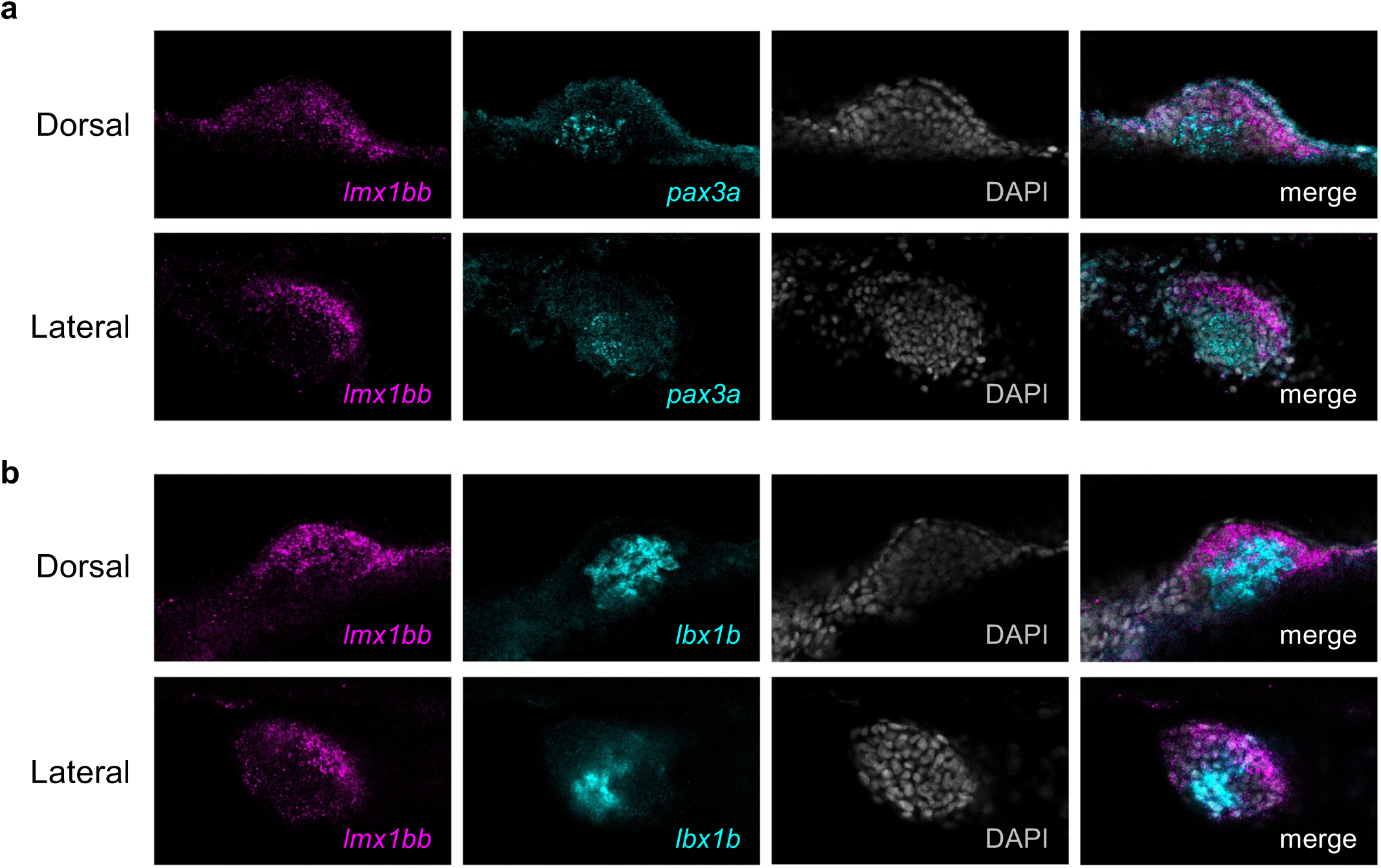
Mesenchymal *lmx1bb*-positive cells form a distinct population from muscle progenitors in pectoral fin buds. HCR co-labelling of *lmx1bb* transcripts with the muscle markers *pax3a* (**a**) and *lbx1b* (**b**) indicates that muscle progenitor populations do not express *lmx1bb* in the fin.

**Supplementary Fig. 2.**
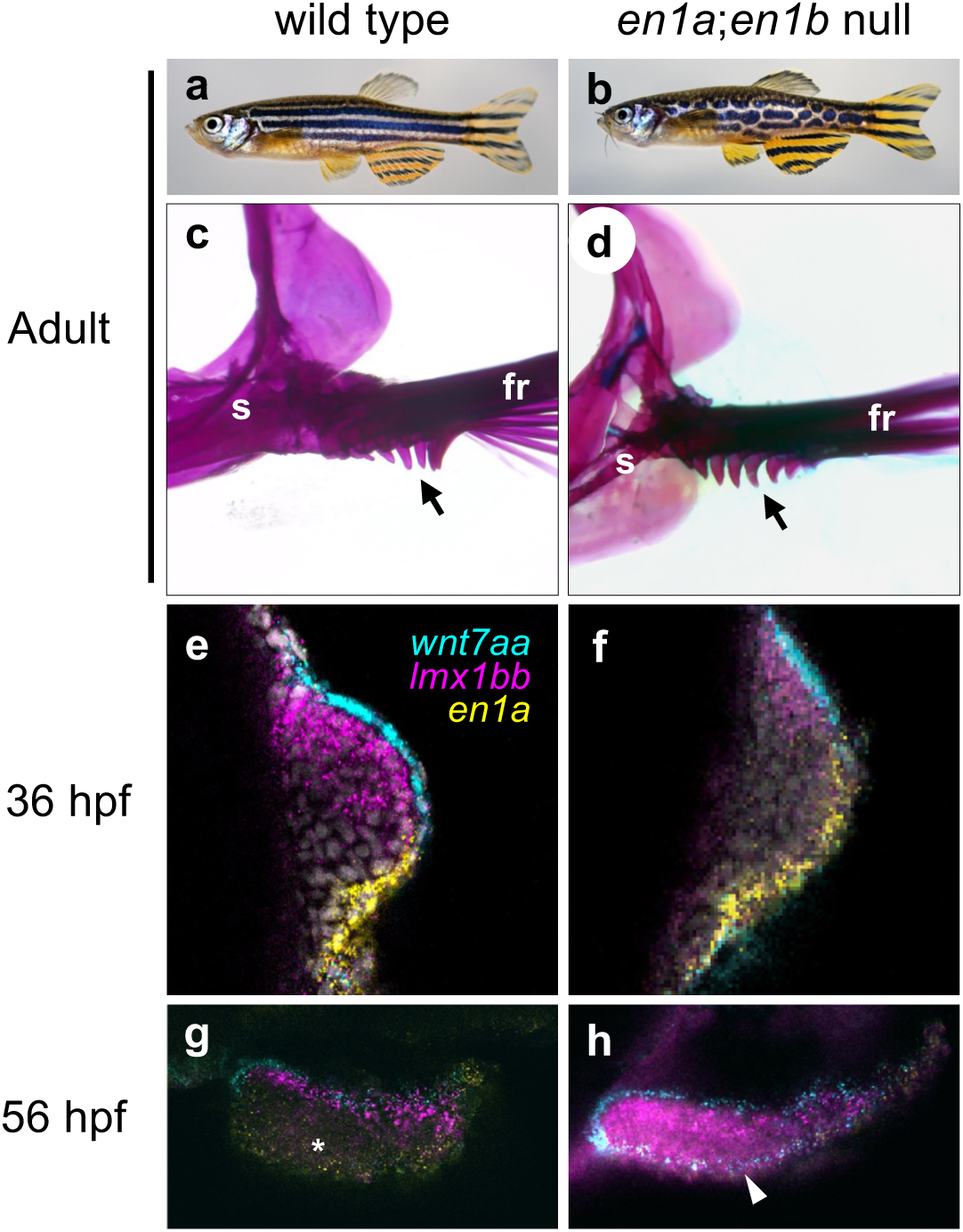
Zebrafish *en1a*;*en1b* double mutants reveal the genetic mechanism of dorsal *Lmx1b* restriction is conserved across bony fishes. **a**, Wildtype adult zebrafish have stipes of pigmentation along the primary body axis (n=10). **b**, *en1a*;*en1b* double mutants exhibit a disrupted pigmentation pattern (n=10). **c**, Wild-type fish develop fin flanges on the ventral surface (black arrow). **d**, The skeleton of double mutant fins retain the ventral flanges, suggesting that En1 function is not required for ventral identity in the proximal fin (n=10). **e**, Expression of DV patterning genes in wild type 36 hpf fin buds. **f**, Double mutants express DV patterning genes in a pattern similar to wild type at 36 hpf (n=5, 100%). **g**, At 56 hpf, the fin has grown outward and *lmx1bb* expression continues to be restricted to the dorsal mesenchyme and the ventral mesenchyme is *lmx1bb* negative (white asterisk, n=5, 100%). **h**, In double mutants, *lmx1bb* is de-repressed and expression is expanded into the ventral mesenchyme by 56 hpf (white arrowhead, n=5, 100%). Anterior to left, dorsal to top in a,b; dorsal to top, distal to left in c-h. fr, fin rays, s, shoulder.

**Supplementary Fig. 3.**
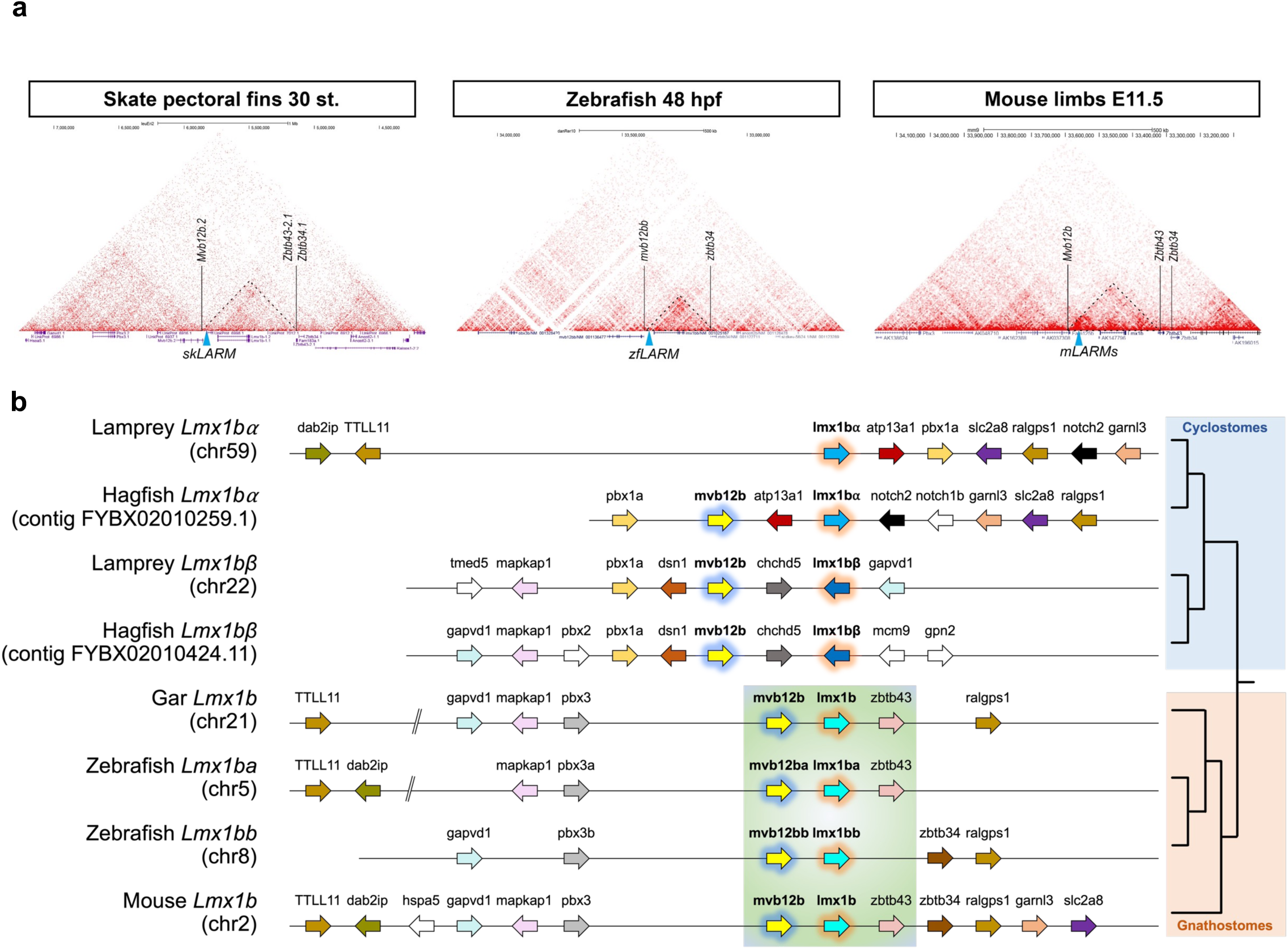
*Mvb12b*-*Lmx1b* synteny is preserved across gnathostomes. **a**, 3D Chromatin confirmation plots reveal the topological association domain containing the *LARM* and *Lmx1b* orthologs is maintained across jawed vertebrates. Hi-C experiments from mouse limbs (Kraft et al., 2019), skate pectoral fins (Marlétaz et al., 2023), and zebrafish embryos (Franke et al., 2021). **b**, Synteny plots demonstrate conservation of the *Mvb12b-Lmx1b* locus among jawed vertebrates. Lmx1b paralogs in cyclostomes (hagfish and lamprey) reside next to similar neighboring genes but without a comparable *Mvb12b-Lmx1b* intergenic region in which the *LARM* elements reside.

**Supplementary Fig. 4.**
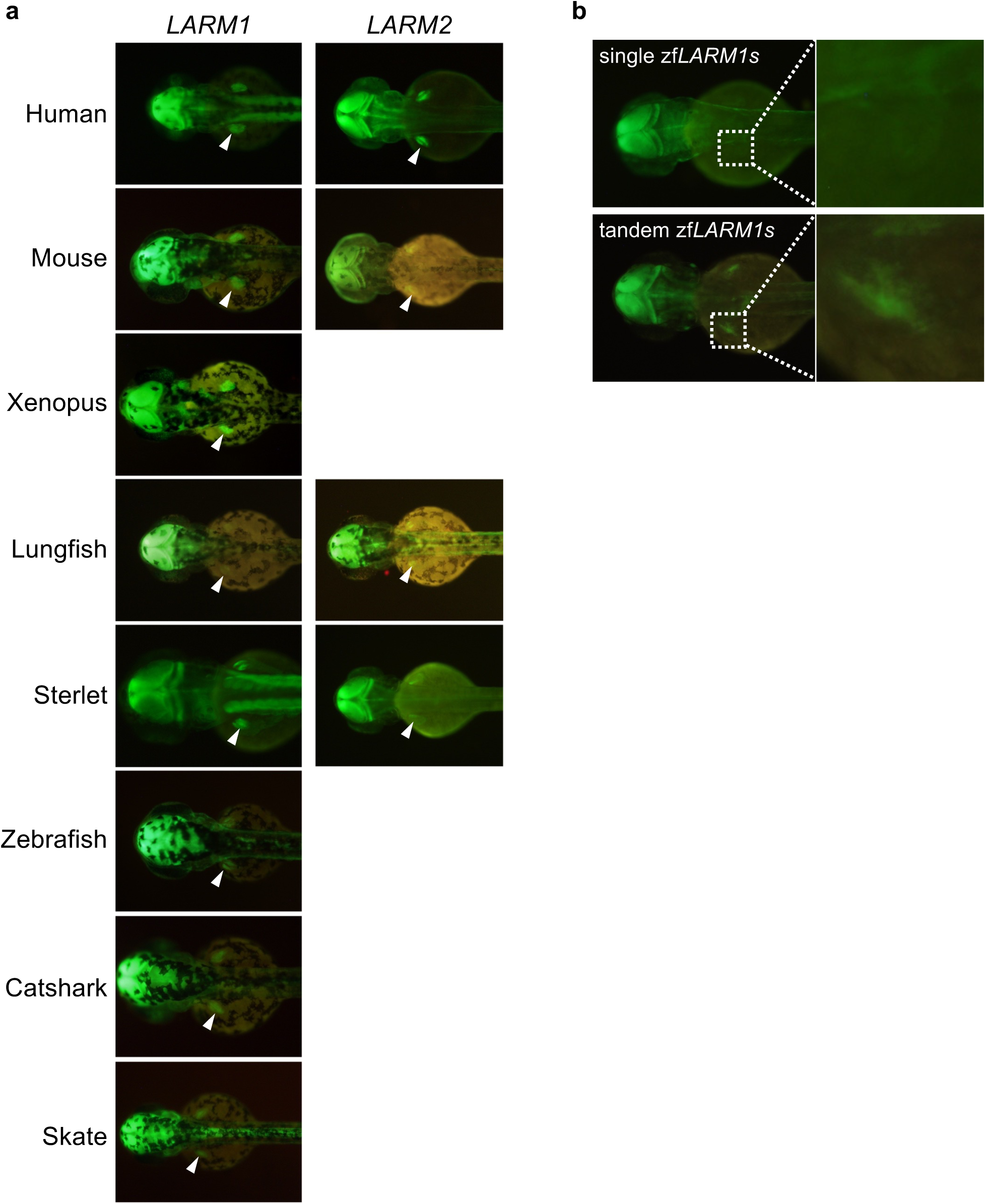
*LARM* elements from across gnathostome evolution drive reporter transgene expression in developing zebrafish pectoral fins. **a**, Reporter expression in F_1_ transgenic animals demonstrates the universal ability of gnathostome *LARM* elements to drive expression in the developing pectoral fins of zebrafish (white arrowhead). **b**, F_1_ transgenic fish carrying a reporter construct with a single *LARM1s* element show limited fin signal, but including a second *LARM1s* in tandem drives elevated reporter expression.

**Supplementary Fig. 5.**
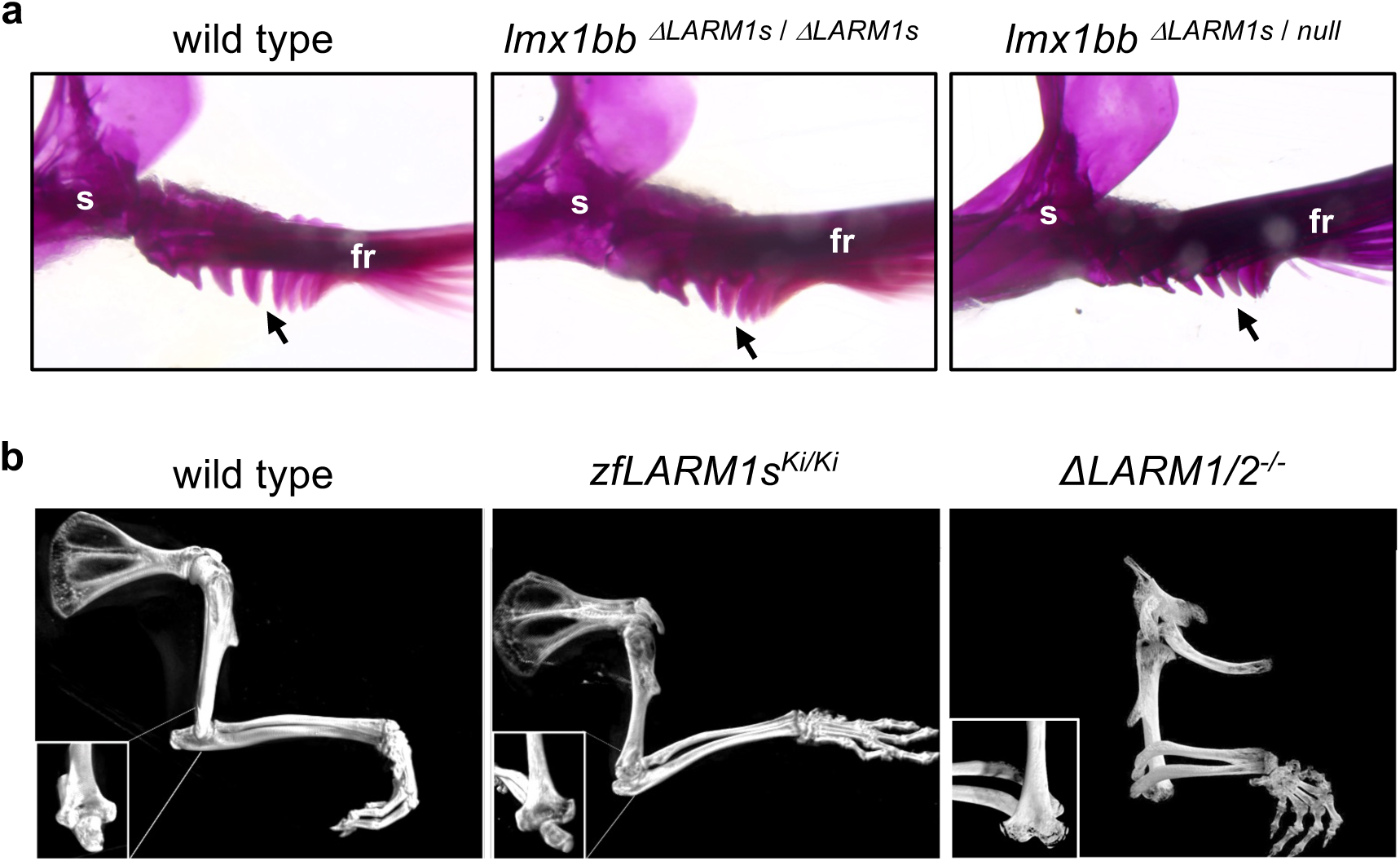
The zebrafish *LARM1s* element is neither necessary nor sufficient for dorsal specification in paired appendages. **a**, Zebrafish homozygous for a deletion of the *LARM1s* element (n=10), or that carry this deletion over a null coding allele (n=10), exhibit wild type fin patterning with ventral flanges only (black arrow). **b**, Mice carrying the zebrafish *LARM1s* element in place of the endogenous murine *LARM* region display partial rescue of the elbow joint dislocation phenotype observed in full *LARM* deletion mice.

**Supplementary Fig. 6.**
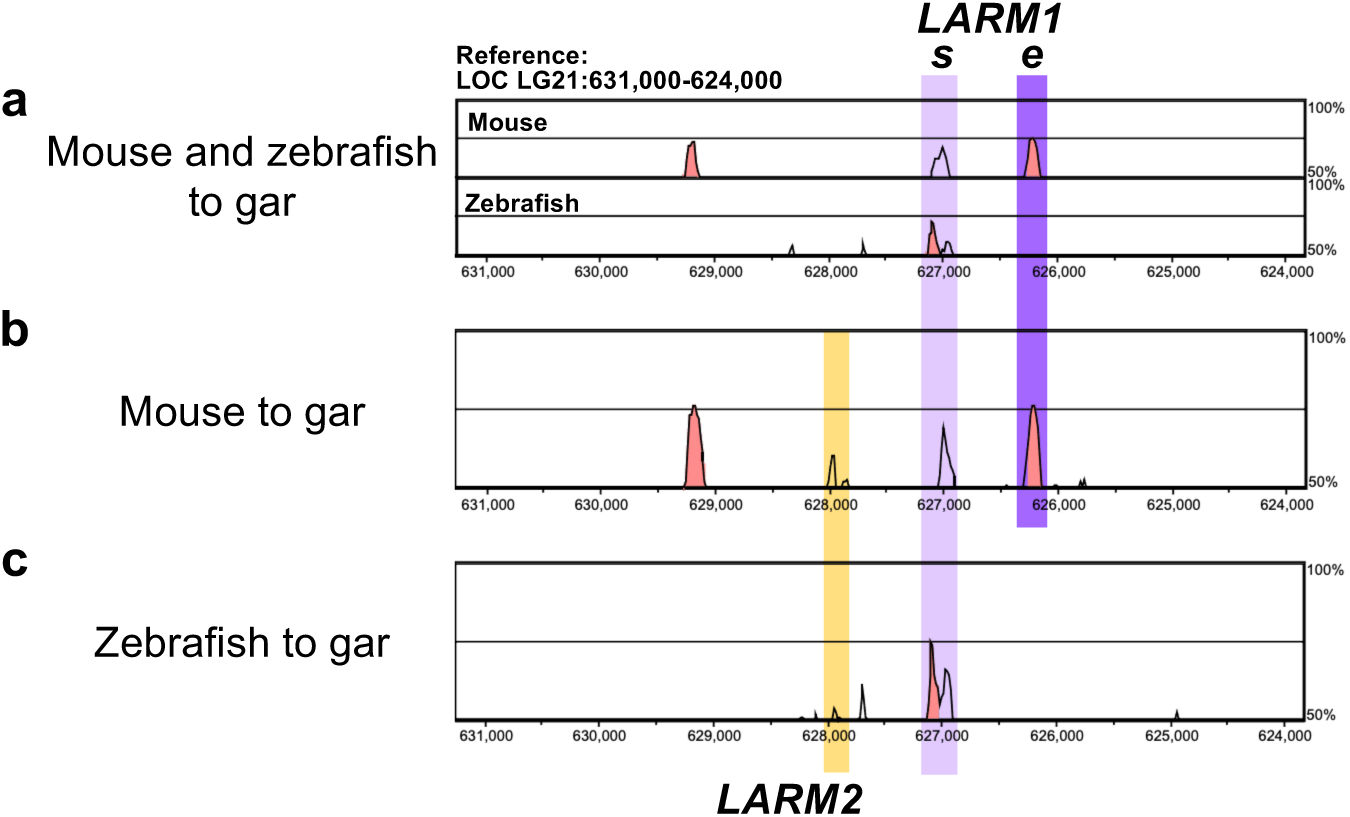
Genomic alignments using the holostean bridge approach identify *LARM1* and *LARM2* elements in the zebrafish. **a**, Plot of a 3-way VISTA alignment of the *Mvb12b-Lmx1b* intergenic region between gar, mouse, and zebrafish. This alignment identifies the *LARM1s* region in all three species (light purple bar), but a peak corresponding to *LARM1e* is only recovered in gar and mouse (dark purple bar) and no *LARM2* peak is detected. **b**, Pairwise alignment of gar and mouse reveals a *LARM2* conservation peak (yellow box) in addition to the *LARM1* regions. **c**, Pairwise alignment of gar and zebrafish identifies a limited *LARM2* conservation peak, but no *LARM1e* region.

**Supplementary Fig. 7.**
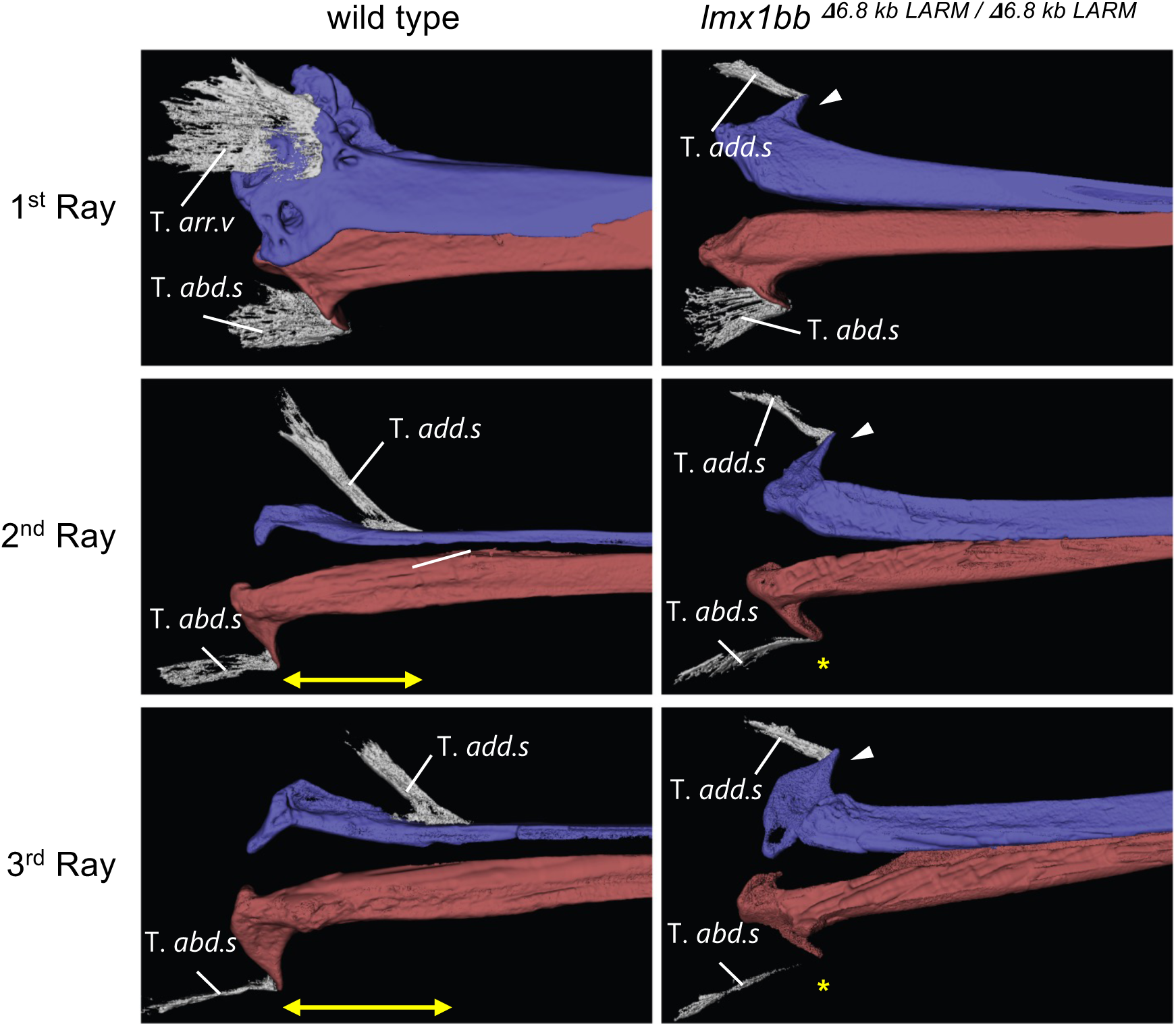
Ventralization of musculoskeletal elements in the dorsal region of holo-*LARM* deletion mutant pectoral fins. Segmented confocal scans of immunolabelled fins reveals full mirror duplication of the ventral configuration in mutant fins. In the 2^nd^ and 3^rd^ rays of wild-type fins, the tendon of the adductor superficialis attaches on the dorsal hemiray distal to the insertion of the abductor superficialis on the ventral hemiray (yellow arrow). In *LARM ^6.8kbΔ/6.8kbΔ^* mutants, the dorsal hemiray develops an ectopic flange (white arrowhead) to which the tendon connects more proximally at a similar level to the insertion on the ventral aspect (yellow asterisk). T. *adb.s*, tendon of abductor superficialis; T. *add.s*, tendon of adductor superficialis; T. *arr.v*, tendon of arrector ventralis.

**Supplementary Fig. 8.**
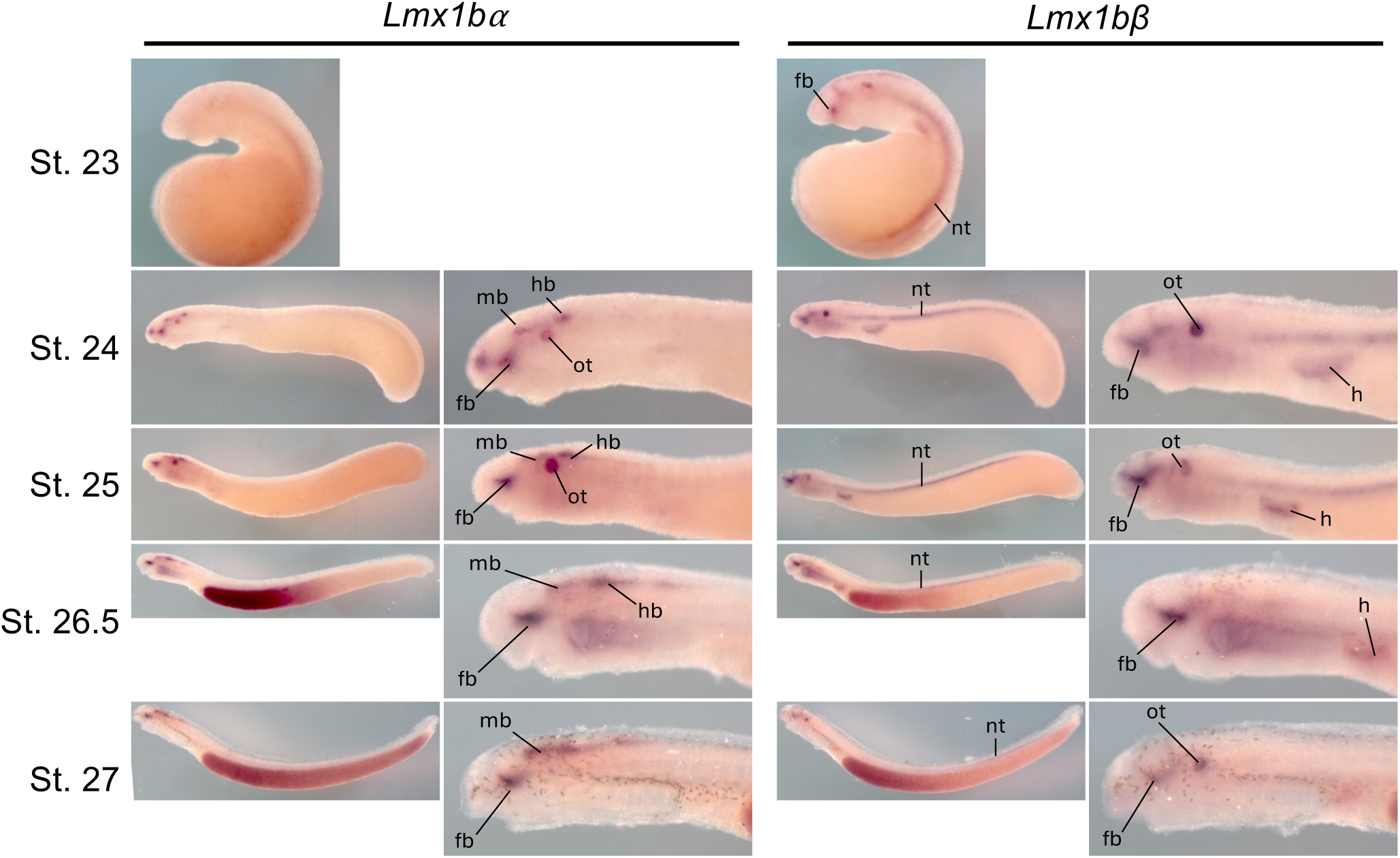
Lamprey *Lmx1b* orthologues are expressed in embryonic domains conserved with gnathostomes. Whole-mount colorometric *in situ* hybridization showing the expression of lamprey *Lmx1b* orthologues. The domains are similar the central nervous system and otic vesicle expression observed in jawed vertebrates. However, *Lmx1b*β is also detected in the heart, an expression domain which is not conserved with gnathostomes. fb, forebrain; h, heart; nt, neural tube; ot, otic vesicle.

**Supplementary Fig. 9.**
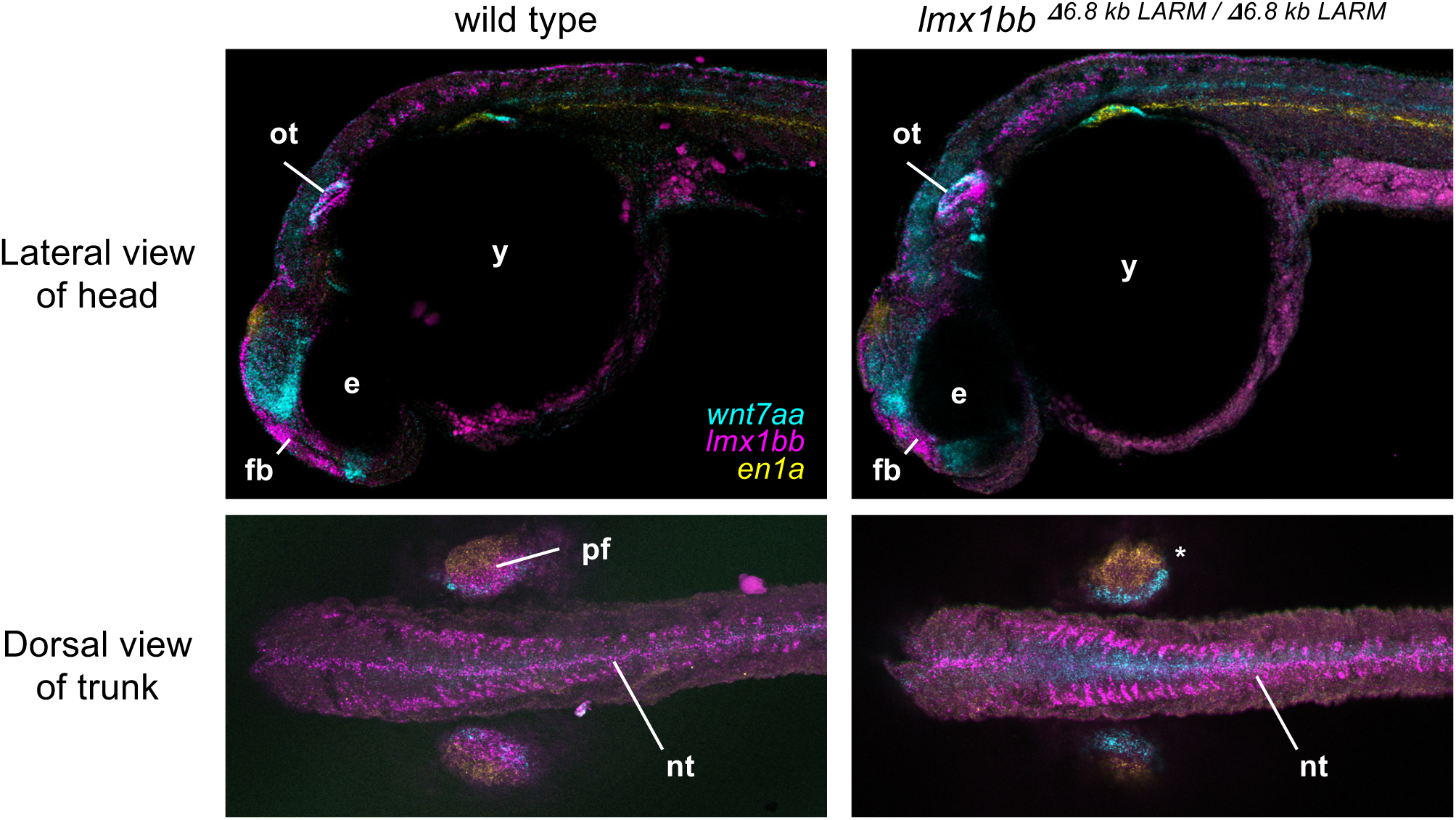
Expression of *lmx1bb* in domains outside of the paired fins is unaffected in holo-*LARM* deletion embryos. HCR co-labelling of *wnt7aa*, *lmx1bb*, and *en1a* transcripts in wild type and *LARM ^6.8kbΔ/6.8kbΔ^* mutant embryos at 36 hpf. Mutant embryos express *lmx1bb* in the same domains as wild type embryos, with the exception of the fin bud (white asterisk). e, eye; fb, forebrain; nt, neural tube; ot, otic vesicle; pf, pectoral fin; y, yolk.

**Supplementary Fig. 10.**
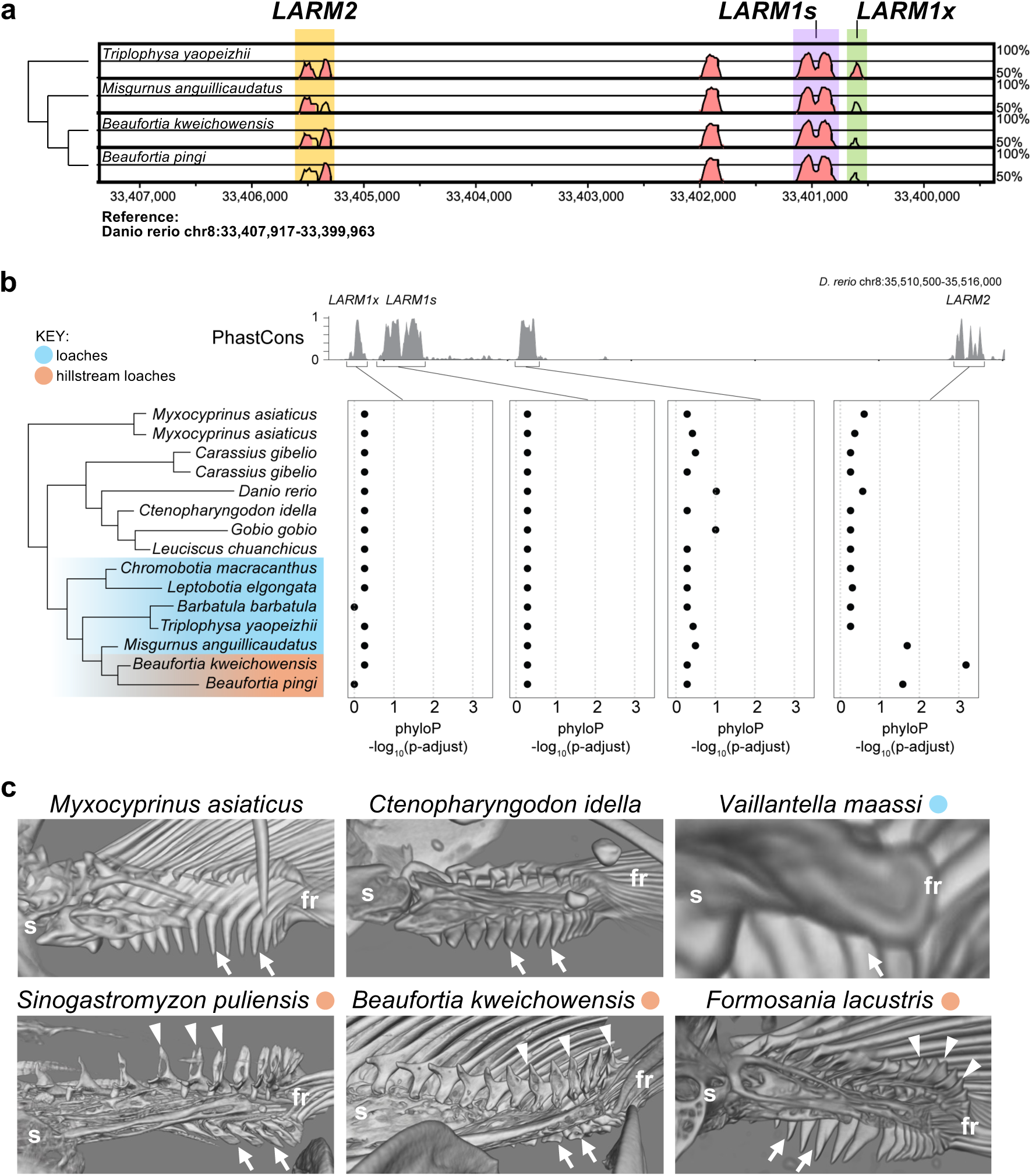
Otherwise conserved *LARM* elements in cypriniform fishes show accelerated sequence evolution in loaches with dorsal pectoral fin elaboration. **a**, VISTA plot of a 5-way multiple sequence alignment between zebrafish and four loach species. The *LARM1s* is highly conserved across the cypriniform order along with *LARM2*. The novel *LARM1x* element is conserved across all non-loach cypriniforms, as well as loach species like *Triplophysa yaopeizhii* outside of the loach families Cobitidate, Balitoridae, and Gastromyzontidae. However, marked reduction in *LARM1x* conservation is observed in cobitid and hillstream loaches and is associated with dorsal pectoral fin elaboration. **b**, Erosion of *LARM1x* in this lineage is due to deletions which are treated as missing data (and not scored) in sequence evolution rate packages such as PhastCons, but an acceleration is detected in the *LARM2* element of these species. **c**, Medial views of cypriniform posterior pectoral fin regions reconstructed from microCT data. Most cypriniform species express a fin configuration similar to zebrafish, with large ventral fin ray flanges (white arrow) and minimal dorsal embellishments (top row). From this ancestral condition, hillstream loaches (bottom row) have evolved dorsal elaboration in the form of dorsal flanges in association with their unique locomotion strategies.

## MATERIALS AND METHODS

### Zebrafish husbandry

Wild-type and genetically modified zebrafish (mutants and transgenics) were maintained and bred under standard conditions (Aleström et al., 2019). Embryos were staged in hours and days post-fertilization (hpf and dpf, respectively) as previously described (Kimmel et al., 1995).

### Hybridization chain reaction (HCR^TM^ RNA-FISH)

Zebrafish: HCR RNA-FISH technique was performed in zebrafish following the official protocol by Molecular Instruments (https://files.molecularinstruments.com/MI-Protocol-RNAFISH-Zebrafish-Rev11.pdf) and Choi et al., (2016), for whole-mount embryos and larvae. Zebrafish embryos were collected at the desired stages, fixed in 4% paraformaldehyde in phosphate buffered saline (PBS) overnight at 4°C and then washed with PBS to stop the fixation. Subsequently, they were dehydrated through an increasing series of methanol (MeOH) solutions in PBS, for their final storage in 100% MeOH at -20°C. Before use, embryos were rehydrated in an inverse series of MeOH/PBST. Juveniles at 14 dpf were permeabilized by incubating them for 7 minutes in a solution containing 10 µg of proteinase K in PBS. This step was skipped for embryos at 31, 36, 48 and 56 hpf. Then they were incubated in a pre-hybridization buffer at 37°C, followed by the addition of specific pairs of DNA probes and overnight incubation at same temperature. Afterwards, washes were performed to remove unbound probes, and the fluorescent signal was triggered and amplified using fluorophore-associated pair of hairpins. Finally, the embryos were washed to remove the excess of hairpins and stained with DAPI to be visualized in a confocal microscope Leica Stellaris 5. Buffers and hairpins were ordered to Molecular Instruments, Inc. DNA probes targeting *lmx1bb*, *wnt7aa*, *en1a*, *lbx1b*, and *pax3a* were designed according to Elagoz et al. (2022), synthesized as oPools (IDT), and used in combination with hairpin amplifiers from Molecular Instruments. The sequences used for probe design are provided in **Supplementary Table 4**.

Mouse: Detection of *Lmx1b* and *En1* transcripts in embryonic mouse limbs was performed following the “Optimized protocol for whole-mount RNA fluorescent in situ hybridization using oxidation-mediated autofluorescence reduction on mouse embryos” described by Morabito et al. (2023). Probe sets targeting *Lmx1b* (NM_010725.3) and *En1* (NM_010133.2) were obtained from Molecular Instruments, along with corresponding B3 and B4 hairpin amplifiers conjugated to Alexa Fluor 647 and Alexa Fluor 488, respectively. Fluorescent images were acquired using a Leica SP5 confocal microscope.

### Allele generation in zebrafish with CRISPR-Cas9

Null alleles of *en1a*, *en1b*, *lmx1ba*, and *lmx1bb* were generated by inducing start codon-deleting mutations or frameshift lesions early in the coding region of each gene using CRISPR-Cas9. Guides were designed using CHOPCHOP (Montague et al., 2014; Labun et al., 2016) and synthesized as crRNAs and used with the Alt-R platform (IDT). Gene-specific crRNAs were duplexed with tracrRNA to a final concentration of 6.25 μM with 1 μg Alt-R^TM^ S.p. Cas9 Nuclease V3 (IDT). This mix was microinjected into T5D strain zebrafish embryos at the one-cell stage. Targeting efficiency was estimated using the T7 endonuclease assay and injected embryos were raised to adulthood (Sentmanat et al., 2018). Injected individuals were outcrossed to wild type animals to recover germline loss-of-function alleles. Genotyping primer sequences are reported in **Supplementary Table 5**, guide sequences are provided in **Supplementary Table 6**, and mutant line information is given in **Supplementary Table 1**.

Enhancer deletion lines removing the zebrafish *LARM1s* element (*LARM^1sΔ^*) or complete *LARM* region (*LARM^6.8^ ^kbΔ^*) were designed using pairs of CRISPR-Cas9 guides flanking the targeted interval to be removed. CHOPCHOP was used to design guides in the zebrafish *LARM* region (Montague et al., 2014; Labun et al., 2016, **Supplementary Table 6**). RNP complexes for each individual guide were assembled as described above and mixed at a 1:1 ratio with their mate guide and microinjected as pairs into single cell T5D zebrafish embryos at a final concentration of 6.25 μM with 1 μg Alt-R^TM^ S.p. Cas9 Nuclease V3 (IDT). Injected clutches were screened by deletion-spanning PCR to detect successful removal. Injected embryos were raised to adulthood and outcrossed to wild type fish, and deletion-spanning PCR was used to identify carriers of the desired deletion. The recovered allele of the *LARM^1sΔ^* deletion carries a clean 188 bp deletion of the targeted element, removing the interval chr8:33401833-33402020 (GRCz11). The recovered *LARM^6.8^ ^kbΔ^* allele removes the 6,845 bp interval between chr8:33400177-33407021 (GRCz11) and replaces it with the 8 bp scar ‘ACTTACTT,’ a motif found within the zfLARM_g4 sequence. Genotyping of the *LARM^1sΔ^* allele was performed with the primers zfLARM1s_F and zfLARM1s_R that yield a 374 bp product from wild type and a 186 bp band in the mutant. The *LARM^6.8^ ^kbΔ^* allele was genotyped using primers zfHoloLARM_F and zfHoloLARM_R, which generate a 7020 bp band in wild type and a 183 bp band in the mutant. Rather than run a PCR to generate a 7020 bp product, *LARM^6.8kbΔ^* incrosses were genotyped with the HoloLARM primers as well as zfLARM primers, which are internal to the large deletion. Animals that showed products from both reactions were considered heterozygous, while animals with a single positive reaction were considered homozygous wild type or homozygous deletion, respective of the positive reaction. Genotyping PCR primers are listed in **Supplementary Table 5**, and deletion line information is given in **Supplementary Table 1**.

The phenotypes reported for each stable mutant zebrafish line are based on a minimum of 10 individuals are were consistent within each genotypic class.

### Mouse strains

C57BL/6J wildtype mice, Δ*LARM1/2* (Haro et al., 2021)*, zfLARM,* and *zfLARM-355* mutant mouse lines were used in this study. All strains were maintained in a C57BL/6J background. The *zfLARM and zfLARM-355* mutants were generated using CRISPR-Cas9 mediated genome editing. To replace the mouse *LARM* region with the zebrafish *LARM1s* ortholog, we used the two gRNAs (355 and 372) described in (Haro et al., 2021; **Supplementary Table 6**) which flank the *LARM1* and *LARM2* mouse elements (chr2:33699834-33707488, mm10). This regulatory region of the *Lmx1b* gene was replaced by homology directed repair (HDR) using a 500-nt single-stranded DNA (ssDNA) donor containing the 164-nt zebrafish *LARM1s* enhancer flanked by 168-nt homology arms corresponding to the 5′ and 3′ deletion breakpoints. The donor sequence was synthesized by GenScript® (GenExactTM ssDNA) and is provided in **Supplementary Table 7**. The zfLARM-355 transgenic line resulted from an unintended CRISPR-Cas9 event, leading to the insertion of two zebrafish *LARM1s* elements in opposite orientations at the 355-gRNA target site. All primers used for genotyping the mutant lines are listed in **Supplementary Table 5**. To genotype the Δ*LARM1/2* deletion, PCR with the 335_Fwd and 335_Rev primers produces a 361 bp product from the wild-type allele, and PCR with the 372_Far_Fwd and 355_Far_Rev primers yields a ∼1.5 kb band in the deletion allele. Genotyping the *zfLARM1s^Ki^* allele was performed with the 355_Fwd and 355_Rev primers that amplify a 361 bp band from the wild-type allele and a 553 bp band from the knockin allele. To detect the *355-zfLARM^Ki^*allele, PCR with the ZF_Fwd and 355_Far_Rev primers will make a 930 bp band in the mutants.

### Skeletal staining

Zebrafish: Adult zebrafish were euthanized by tricaine overdose and fixed overnight in 3.7% formaldehyde in phosphate buffered saline (PBS) at room temperature with agitation. After fixation, specimens were washed in PBS for one hour, then dehydrated through a stepped series of 2-hour washes of graded ethanol series into 100% ethanol. Following dehydration, animals were stained in Alcian Blue (70% ethanol, 30% acetic acid, 0.015% Alcian Blue) overnight at room temperature with rocking. After staining specimens were washed in 95% ethanol for 1 hour and then rehydrated through a stepped series of 2-hour washes of ethanol, and then washed in tap water twice for 1 hour each. Rehydrated specimens were then macerated in trypsin solution (0.3% bovine trypsin powder in 65% saturated sodium borax solution) for 90 minutes at 37°C. Specimens were then stained overnight in Alizarin solution (0.25% in 0.5% potassium hydroxide (KOH) solution) at room temperature with agitation. Specimens were washed in 0.5% KOH for 1 day and then moved into 100% glycerol.

Mouse: Limbs from 1-month-old mice were collected, skinned, and fixed overnight (ON) in 95% ethanol. For cartilage staining, samples were incubated in Alcian Blue solution (80% ethanol, 20% glacial acetic acid, and 0.3 mg/ml Alcian Blue) for 48–72 hours, then rinsed twice in 95% ethanol for 1–2 days. Limbs were subsequently cleared in 1% potassium hydroxide (KOH) for 24 hours or until the skeletal elements became visible. Bone staining was performed by incubating samples in 0.05 mg/ml Alizarin Red in 1% KOH, in the dark, for 24–48 hours. Following staining, samples were cleared in 20% glycerol/1% KOH for 24 hours and then dehydrated through a graded series of 70% ethanol:glycerol:water mixtures (1:2:7; 3:3:4; 4:4:2; 5:5:0). Finally, skeletal preparations were photographed and stored in 70% ethanol:glycerol:water (5:5:0).

### Analysis of Hi-C data

To analyze the conservation of the *Lmx1b* 3D genome architecture we used publicly available Hi-C data from pectoral fins of stage 30 skate embryos (Marlétaz et al., 2023), 48 hpf zebrafish embryos (Franke et al., 2021), and limb buds of E11.5 mice (Kraft et al., 2019), The chromatin contacts within the *Mvb12b-Zbtb34/43* locus were visualized in the UCSC Genome Browser (http://genome.ucsc.edu; Perez et al., 2025).

### Identification of *LARM* orthologue sequences in fishes

We generated a Hidden Markov Model (HMM) for deep sequence homology using ‘nhmmer’ (Wheeler & Eddy, 2013) to look for homologous regions in vertebrate genomes where we knew the *Lmx1b* locus was strongly conserved (chicken, turkey, coelacanth, human, bat, whale, cow, manatee, otter, seal, gecko, and mouse). The *LARM* sequences of the mentioned species were retrieved using a blastn with the mouse *LARM* elements (*LARM1s* chr2:33701467-33701677; *LARM1e* chr2:33700794-33701108; *LARM2*: chr2:33705926-33706432; mm10 coordinates). Then, we performed a Multiple Sequence Alignment (MSA) using ‘clustalw2’ (Larkin et al., 2007) with the default parameters for DNA sequences, and this MSA was used to generate an HMM profile using ‘hmbuild’ from HMMER (Wheeler & Eddy, 2013). The database for homology searching consists in a multi FASTA file containing the genomic region around the *Lmx1b* gene spanning 30 times the length of their annotated *Lmx1b* of different fish species (skate, catshark, spotted gar, sterlet sturgeon, zebrafish, medaka, coelacanth, and Australian lungfish). This length was arbitrary defined with the idea of covering the *Lmx1b* regulatory landscape where the *LARM* elements are contained.

### Enhancer reporter assays in zebrafish

Transgenic embryos were generated using the Tol2 transposase system (Kawakami, 2007). The sequences of assayed *LARM* elements are provided in **Supplementary Table 2**, and the primers used to amplify them are given in **Supplementary Table 5**. Putative *LARM* regulatory regions were amplified from genomic DNA by PCR, except for lungfish and sterlet elements, as well as the zebrafish *LARM1s*, which were synthetized by Genewiz (Azenta Life Sciences). PCR products were purified with the Isolate II PCR and Gel Kit (BIOLINE) and cloned into an intermediate vector (pCR8/GW/TOPO, Invitrogen, #10532893). Using Gateway® technology (LR Clonase® II Enzyme mix, Invitrogen, ThermoFisher Scientific), amplicons were finally recombined into an enhancer detection vector containing the *gata2* minimal promoter and the strong midbrain *irx* enhancer *z48* for an internal control of the transgenesis experiment (Kawakami et al., 2004; Gehrke et al., 2015). Briefly, one-cell stage embryos were microinjected with 2-3 nl of a solution containing 30 ng/μl of transposase mRNA, 20 ng/μl of purified enhancer construct, and 0.05% phenol red solution. The F_0_ injected embryos were monitored at 24, 48, and 72 hpf for GFP expression (midbrain control and enhancer-specific). GFP+ embryos with enhancer activity in the pectoral fins were raised to sexual maturity and crossed with wildtype animals to obtain F_1_ animals. The F_1_ with fin enhancer activity were crossed with wildtype animals, raised and the F_2_ descendants carrying the enhancer construct were considered as stable transgenic lines. An enhancer was considered to have verified activity in pectoral fins when three or more independent stable transgenic lines from the same construct exhibit similar and consistent expression patterns in the fin. Transgenic embryos were imaged using an Olympus SZX16 fluorescence stereoscope and photographed with an Olympus DP71 camera.

### Whole-mount colorometric *in situ* hybridization

Zebrafish: The detection of reporter-derived GFP transcripts was performed according to (Jowett & Lettice, 1994). Briefly, GFP transgenic zebrafish embryos were fixed at the appropriate developmental stage and dehydrated in a Methanol/PBS increasing series. Embryos were rehydrated in a reverse Methanol/PBS-Tween series, prehybridized in hybridization buffer and finally hybridized overnight with 2 ng/µl of the GFP digoxigenin-labeled RNA probe at 70°C. The excess of probe was washed with a decreasing formamide buffers series to reduce the unbound probe background. Embryos were blocked and incubated with the anti-digoxigenin-Alkaline Phosphatase (AP) conjugated antibody solution. AP activity was finally revealed using a staining solution of NBT-BCIP.

Mouse: Mouse embryos were fixed in PFA 4% ON at 4°C, washed in PBS, in PBT (0.1% Tween-20 in PBS), and dehydrated with MeOH by increased concentrations until 100% MeOH. Next day, samples were rehydrated, washed in PBT and bleached with 6% H2O2/PBT for 1h. Embryos were washed in PBT and digested with Proteinase K (PK, Roche, #03115879001) at 10 μg/ml in PK buffer (50 mM Tris-HCl pH 7.4, 5 mM EDTA), adjusting the digestion time to the embryonic stage. Then, embryos were quickly washed in PBT and post-fixed in 0.25% glutaraldehyde in 4% PFA during 20 min, and leaved in the hybridization buffer at 65°C ON (50% formamide, 5x SSC (saline-sodium citrate), 2% blocking powder, 0.1% Triton X-100, 0.1% CHAPS, 1 mg/ml tRNA, 50 μg/ml Heparin pH 4.5, 500 mM EDTA pH 8). The following day, embryonic samples were frozen at -20°C for at least 6 h, and later incubated with the hybridization buffer containing the desired antisense RNA probe. The next day, several post-hybridization washes (65°C) were carried out to remove nonspecific binding: 3X 2xSSC/0.1% CHAPS 30 min each, and 3X 0.2xSSC/0.1% CHAPS (20-20-30 min). Then, samples were washed twice with KTBT buffer (50 mM Tris-HCl pH 7.4, 150 mM NaCl, 10 mM KCl, 1%Triton X-100) for 10 min at RT, and later blocked in 20% sheep serum/KTBT for 2 h at RT. Embryos were incubated with Anti-Digoxigenin-AP (Roche, #11093274910) diluted 1:2,000 in blocking solution at 4°C ON. After this, several washes were performed with KTBT at RT. Finally, signal detection was performed by incubating the embryos in darkness in NTMT buffer (100 mM Tris-HCl pH 9.5, 50 mM MgCl2, 100 mM NaCl, 1% Triton X-100), with NBT (3 μg/ml) and BCIP (2.3 μg/ml) substrates. Once the desired signal level was reached, reaction was stopped with several washes in KTBT and fixed in 4% PFA for photographing.

Lamprey: Two *Lmx1b* genes are annotated in the kPetMar1.pri Feb. 2020 sea lamprey genome assembly (GCF_010993605.1), one on chromosome 59 (XM_032976538.1) which we have called *Lmx1bα* and the other on chromosome 22 (XM_032959033.1) which we have called *Lmx1bβ*. To generate *in situ* hybridization probe templates, we ordered from GENEWIZ/Azenta synthesized DNA fragments corresponding to exon-spanning segments of either transcript flanked by Sp6, T7, and M13 primer sites (**Supplementary Table 8**). These templates were used to synthesize digoxigenin-labelled RNA probes as previously described (Lufkin, 2007). Embryonic and larval material was obtained as in Cattell et al., (2011) and animals were staged according to Tahara (1988). Whole-mount *in situ* hybridization was performed following Meulemans and Bronner-Fraser (2002) with the modification that proteinase K treatment for 6 cm ammocete larvae for 20 minutes at a concentration of 1.6 IU/mL in PBST.

### Micro computed tomography (**μ**CT)

Limbs from 1-month-old mice were scanned using a Skyscan 1172 micro-CT scanner (Bruker) at 40 kV, 100 μA, and a pixel resolution of 27.0 μm. Image reconstruction was performed using NRecon software (Bruker), and 3D renderings were generated with CTvox v3.3.1 (Bruker).

### Immunolabelling of pectoral fin musculoskeletal elements

Immunolabelling was performed using a skeletal muscle antibody (12/101, Developmental Studies Hybridoma Bank) and anti-thrombospondin 4 antibody (Abcam). The labelling protocol followed the Deepclear method (Pende et al. 2020), with several modifications, including an overnight clearing in solution 1.1, primary antibody incubation for 3 days at 37°C, incubation in the secondary for two days at room temperature, and a five-minute incubation in alizarin red S for bone counterstaining, with a final clearing step using methanol dehydration, followed by dichloromethane incubation and methyl cinnamate. Images were taken at 20x using a Leica Stellaris confocal microscope.

### VISTA genome alignments

To identify conserved *LARM* elements, the *Mvb12b-Lmx1b* intergenic region from different species were aligned using mVISTA (https://genome.lbl.gov/vista/index.shtml, Dubchak et al., 2000; Frazer et al., 2004). Sequences were aligned using default parameters and the LAGAN alignment program (Brundo et al., 2003).

### 3D Reconstruction of cypriniform pectoral fin skeletons

All cypriniform computed microCT data sets were accessed on MorphoSource (www.morphosource.org; Blackburn et al., 2024). The ARK identifiers and specimen information are listed in **Supplementary Table 9**. Access to these open source data was provided with support from oVert TCN, NSF DBI-1701753, and NSF DBI-1701714. Reconstructions were down sampled to 8-bit and cropped to the pectoral fin region of interest using ImageJ (Schindelin et al., 2015). Processed reconstructions were visualized and imaged in Amira 6 (FEI Systems).

### Analysis of *LARM1x* element in hillstream loaches

To verify the *LARM1x* deletions in the *Beaufortia* genomic data, we cloned this region from *Sewellia lineolata* using using PCR with the primers LARM1x_F and LARM1x_R (**Supplementary Table 10**).

### Evolutionary rate analysis of cypriniform *LARM* elements

To characterize the *LARM* region across order Cypriniformes, we used BLAST+ v2.14.1 (Camacho et al., 2020) to identify sequences orthologous to *LARM1* across a series of cypriniform genomes: *Barbatula barbatula* (GCA_037178815.1), *Beaufortia kweichowensis* (GCA_019155185.1), *Beaufortia pingi* (GCA_044048495.1), Carassius gibelio (GCA_023724105.1), *Chromobotia macracanthus* (GCA_036877525.1), *Ctenopharyngodon idella* (GCF_019924925.1), *Danio rerio* (GCF_049306965.1), *Gobio gobio* (GCA_949357685.1), *Leptobotia elgongata* (GCA_039881065.1), *Leuciscus chuanchicus* (GCA_965140175.1), *Misgurnus anguillicaudatus* (GCF_027580225.2), *Myxocyprinus asiaticus* (GCF_019703515.2), *Triplophysa yaopeizhii* (GCA_048296945.1). For *Carassius gibelio* and *Myxocyprinus asiaticus*, which have undergone recent, independent whole-genome duplications, we included both orthologous *LARM* loci.

Each identified *LARM* locus was extended by 250 kilobases (kb) upstream and downstream to create a 500 kb region. These extended loci were aligned using Progressive Cactus v2.9.9 (Armstrong et al., 2020) to generate a multiple genome alignment. The resulting Hierarchical Alignment (HAL) file was converted to a Multiple Alignment File (MAF) using hal2maf (parameters ‘--chunkSize 10000 --noAncestors -- dupeMode single’).

The cladogram used in the Cactus alignment and all subsequent analyses was based on Stout et al., 2016, with the exception that Wang et al., 2016 was used for the topology within loaches provided its greater sampling of loach lineages. The resulting composite cladogram is:(((((((beaufortia_pingi,beaufortia_kweichowensis),misgurnus_anguillicaudatus),(triplo physa_yaopeizhii,barbatula_barbatula)),(leptobotia_elgongata,chromobotia_macracanth us)),(((leuciscus_chuanchicus,gobio_gobio),ctenopharyngodon_idella),danio_rerio)),(ca rassius_gibelio,carassius_gibelio2)),(myxocyprinus_asiaticus,myxocyprinus_asiaticus2)).

To identify conserved elements within the intergenic region between *lmx1bb* and *mvb12bb*, we applied PhastCons (Siepel et al., 2005; Hubisz et al., 2011) on the 500 kb MAF (PHAST v1.5; parameters ‘--target-coverage 0.25 --expected-length 12 --msa-format MAF). Prior to running PhastCons, we generated a phylogenetic model using phyloFit (PHAST v1.5) based on the full 500 kb extended LARM locus. We then calculated a 20 bp sliding window average (with 1 bp step size) of the PhastCons score across the resulting wiggle (WIG) file to identify peaks of conservation. Within these peaks we identified three discrete conserved *LARM* elements: *LARM1x*, *LARM1s*, and *LARM2*. Within each *LARM*, we evaluated coverage and conservation by comparing the sequences of each species to *Danio rerio*. To test for accelerated sequence evolution in these regions, we used phyloP (Pollard et al., 2010) (PHAST v1.5) on each LARM (parameters ‘--mode ACC --method LRT --features --subtree’).

We assessed the presence or absence of transcription factor binding site motifs across the phylogeny using the Find Individual Motif Occurrences tool (FIMO v5.5.5)(Grant et al., 2011) and the JASPAR2022 CORE non-redundant database (Sandelin et a., 2004). Motifs that were specifically gained or lost within *Beaufortia* (FIMO q-value < 0.05) in the *LARM1x* region were recorded (**Supplementary Table 3**).

## REFERENCES

Aleström P, D’Angelo L, Midtlyng PJ, Schorderet DF, Schulte-Merker S, Sohm F, et al. Zebrafish: Housing and husbandry recommendations. Lab Anim 2020;54:213–24.

Allou L, Balzano S, Magg A, Quinodoz M, Royer-Bertrand B, Schöpflin R, et al. Non-coding deletions identify Maenli lncRNA as a limb-specific En1 regulator. Nature 2021;592:93–8.

Armstrong J, Hickey G, Diekhans M, Fiddes IT, Novak AM, Deran A, et al. Progressive Cactus is a multiple-genome aligner for the thousand-genome era. Nature 2020;587:246–51.

Blackburn DC, Boyer DM, Gray JA, Winchester J, Bates JM, Baumgart SL, et al. Increasing the impact of vertebrate scientific collections through 3D imaging: The openVertebrate (oVert) Thematic Collections Network. Bioscience 2024;74:169–86.

Braasch I, Gehrke AR, Smith JJ, Kawasaki K, Manousaki T, Pasquier J, et al. The spotted gar genome illuminates vertebrate evolution and facilitates human-teleost comparisons. Nat Genet 2016;48:427–37.

Brudno M, Do CB, Cooper GM, Kim MF, Davydov E, NISC Comparative Sequencing Program, et al. LAGAN and Multi-LAGAN: efficient tools for large-scale multiple alignment of genomic DNA. Genome Res 2003;13:721–31.

Burzynski GM, Reed X, Maragh S, Matsui T, McCallion AS. Integration of genomic and functional approaches reveals enhancers at LMX1A and LMX1B. Mol Genet Genomics 2013;288:579–89.

Camacho C, Coulouris G, Avagyan V, Ma N, Papadopoulos J, Bealer K, et al. BLAST+: architecture and applications. BMC Bioinformatics 2009;10:421.

Cattell M, Lai S, Cerny R, Medeiros DM. A new mechanistic scenario for the origin and evolution of vertebrate cartilage. PLoS One 2011;6:e22474.

Chen H, Lun Y, Ovchinnikov D, Kokubo H, Oberg KC, Pepicelli CV, et al. Limb and kidney defects in Lmx1b mutant mice suggest an involvement of LMX1B in human nail patella syndrome. Nat Genet 1998;19:51–5.

Chen H, Ovchinnikov D, Pressman CL, Aulehla A, Lun Y, Johnson RL. Multiple calvarial defects in lmx1b mutant mice. Dev Genet 1998;22:314–20.

Chen XJ, Song L, Liu WZ. Seven fish complete mitochondrial genomes of the Gastromyzontidae in Southern China (Teleostei: Cypriniformes). Mitochondrial DNA B Resour 2022;7:624–6.

Choi HMT, Calvert CR, Husain N, Huss D, Barsi JC, Deverman BE, et al. Mapping a multiplexed zoo of mRNA expression. Development 2016;143:3632–7.

Crawford CH, Randall ZS, Hart PB, Page LM, Chakrabarty P, Suvarnaraksha A, et al. Skeletal and muscular pelvic morphology of hillstream loaches (Cypriniformes: Balitoridae). J Morphol 2020;281:1280–95.

Dahn RD, Davis MC, Pappano WN, Shubin NH. Sonic hedgehog function in chondrichthyan fins and the evolution of appendage patterning. Nature 2007;445:311–4.

Dealy CN, Roth A, Ferrari D, Brown AMC, Kosher RA. Wnt-5a and Wnt-7a are expressed in the developing chick limb bud in a manner suggesting roles in pattern formation along the proximodistal and dorsoventral axes. Mech Dev 1993;43:175–86.

Dubchak I, Brudno M, Loots GG, Pachter L, Mayor C, Rubin EM, et al. Active conservation of noncoding sequences revealed by three-way species comparisons. Genome Res 2000;10:1304–6.

Ekker M, Wegner J, Akimenko MA, Westerfield M. Coordinate embryonic expression of three zebrafish engrailed genes. Development 1992;116:1001–10.

Elagoz AM, Styfhals R, Maccuro S, Masin L, Moons L, Seuntjens E. Optimization of whole mount RNA multiplexed in situ hybridization chain reaction with immunohistochemistry, clearing and imaging to visualize octopus embryonic neurogenesis. Front Physiol 2022;13:882413.

Franke M, De la Calle-Mustienes E, Neto A, Almuedo-Castillo M, Irastorza-Azcarate I, Acemel RD, et al. CTCF knockout in zebrafish induces alterations in regulatory landscapes and developmental gene expression. Nat Commun 2021;12:1–19.

Frazer KA, Pachter L, Poliakov A, Rubin EM, Dubchak I. VISTA: computational tools for comparative genomics. Nucleic Acids Res 2004;32:W273–9.

Freitas R, Zhang G, Cohn MJ. Evidence that mechanisms of fin development evolved in the midline of early vertebrates. Nature 2006;442:1033–7.

Gehrke AR, Schneider I, de la Calle-Mustienes E, Tena JJ, Gomez-Marin C, Chandran M, et al. Deep conservation of wrist and digit enhancers in fish. Proc Natl Acad Sci U S A 2015;112:803–8.

Grandel H, Draper BW, Schulte-Merker S. dackel acts in the ectoderm of the zebrafish pectoral fin bud to maintain AER signaling. Development 2000;127:4169–78.

Grant CE, Bailey TL, Noble WS. FIMO: scanning for occurrences of a given motif. Bioinformatics 2011;27:1017–8.

Haro E, Petit F, Pira CU, Spady CD, Lucas-Toca S, Yorozuya LI, et al. Identification of limb-specific Lmx1b auto-regulatory modules with Nail-patella syndrome pathogenicity. Nat Commun 2021;12:5533.

Haro E, Watson BA, Feenstra JM, Tegeler L, Pira CU, Mohan S, et al. Lmx1b-targeted cis-regulatory modules involved in limb dorsalization. Development 2017;144:2009–20.

Havird JC, Page LM. A revision of Lepidocephalichthys (teleostei: Cobitidae) with descriptions of two new species from Thailand, Laos, Vietnam, and Myanmar. Copeia 2010;2010:137–59.

Hawkins MB, Jandzik D, Tulenko FJ, Cass AN, Nakamura T, Shubin NH, et al. An Fgf-Shh positive feedback loop drives growth in developing unpaired fins. Proc Natl Acad Sci U S A 2022;119:e2120150119.

Hubisz MJ, Pollard KS, Siepel A. PHAST and RPHAST: phylogenetic analysis with space/time models. Brief Bioinform 2011;12:41–51.

Jagla K, Dollé P, Mattei MG, Jagla T, Schuhbaur B, Dretzen G, et al. Mouse Lbx1 and human LBX1 define a novel mammalian homeobox gene family related to the Drosophila lady bird genes. Mech Dev 1995;53:345–56.

Jowett T, Lettice L. Whole-mount in situ hybridizations on zebrafish embryos using a mixture of digoxigenin- and fluorescein-labelled probes. Trends Genet 1994;10:73–4.

Jung H, Baek M, D’Elia KP, Boisvert C, Currie PD, Tay B-H, et al. The ancient origins of neural substrates for land walking. Cell 2018;172:667–682.e15.

Kang J, Nachtrab G, Poss KD. Local Dkk1 crosstalk from breeding ornaments impedes regeneration of injured male zebrafish fins. Dev Cell 2013;27:19–31.

Kawakami K. Tol2: a versatile gene transfer vector in vertebrates. Genome Biol 2007;8 Suppl 1:S7.

Kawakami K, Takeda H, Kawakami N, Kobayashi M, Matsuda N, Mishina M. A transposon-mediated gene trap approach identifies developmentally regulated genes in zebrafish. Dev Cell 2004;7:133–44.

Kimmel CB, Ballard WW, Kimmel SR, Ullmann B, Schilling TF. Stages of embryonic development of the zebrafish. Dev Dyn 1995;203:253–310.

Kraft K, Magg A, Heinrich V, Riemenschneider C, Schöpflin R, Markowski J, et al. Serial genomic inversions induce tissue-specific architectural stripes, gene misexpression and congenital malformations. Nat Cell Biol 2019;21:305–10.

Labun K, Montague TG, Gagnon JA, Thyme SB, Valen E. CHOPCHOP v2: a web tool for the next generation of CRISPR genome engineering. Nucleic Acids Res 2016;44:W272–6.

Labun K, Montague TG, Krause M, Torres Cleuren YN, Tjeldnes H, Valen E. CHOPCHOP v3: expanding the CRISPR web toolbox beyond genome editing. Nucleic Acids Res 2019;47:W171–4.

Larkin MA, Blackshields G, Brown NP, Chenna R, McGettigan PA, McWilliam H, et al. Clustal W and Clustal X version 2.0. Bioinformatics 2007;23:2947–8.

Letelier J, de la Calle-Mustienes E, Pieretti J, Naranjo S, Maeso I, Nakamura T, et al. A conserved Shh cis-regulatory module highlights a common developmental origin of unpaired and paired fins. Nat Genet 2018;50:504–9.

Letelier J, Naranjo S, Sospedra-Arrufat I, Martínez-Morales J, López-Ríos J, Shubin N, et al. The Shh/Gli3 gene regulatory network precedes the origin of paired fins and reveals the deep homology between distal fins and digits. Proc Natl Acad Sci U S A 2020;118:. 10.1073/pnas.2100575118.

Logan C, Hornbruch A, Campbell I, Lumsden A. The role of Engrailed in establishing the dorsoventral axis of the chick limb. Development 1997;124:2317–24.

Loomis CA, Harris E, Michaud J, Wurst W, Hanks M, Joyner AL. The mouse Engrailed-1 gene and ventral limb patterning. Nature 1996;382:360–3.

Lufkin T. In situ hybridization of whole-mount mouse embryos with RNA probes: hybridization, washes, and histochemistry. CSH Protoc 2007;2007:db.prot4823.

Marlétaz F, de la Calle-Mustienes E, Acemel RD, Paliou C, Naranjo S, Martínez-García PM, et al. The little skate genome and the evolutionary emergence of wing-like fins. Nature 2023;616:495–503.

McMillan SC, Xu ZT, Zhang J, Teh C, Korzh V, Trudeau VL, et al. Regeneration of breeding tubercles on zebrafish pectoral fins requires androgens and two waves of revascularization. Development 2013;140:4323–34.

Meulemans D, Bronner-Fraser M. Amphioxus and lamprey AP-2 genes: implications for neural crest evolution and migration patterns. Development 2002;129:4953–62.

Minchin JEN, Williams VC, Hinits Y, Low S, Tandon P, Fan C-M, et al. Oesophageal and sternohyal muscle fibres are novel Pax3-dependent migratory somite derivatives essential for ingestion. Development 2013;140:2972–84.

Montague TG, Cruz JM, Gagnon JA, Church GM, Valen E. CHOPCHOP: a CRISPR/Cas9 and TALEN web tool for genome editing. Nucleic Acids Res 2014;42:W401–7.

Morabito A, Malkmus J, Pancho A, Zuniga A, Zeller R, Sheth R. Optimized protocol for whole-mount RNA fluorescent in situ hybridization using oxidation-mediated autofluorescence reduction on mouse embryos. STAR Protoc 2023;4:102603.

Neumann CJ, Grandel H, Gaffield W, Schulte-Merker S, Nüsslein-Volhard C. Transient establishment of anteroposterior polarity in the zebrafish pectoral fin bud in the absence of sonic hedgehog activity. Development 1999;126:4817–26.

Parr BA, McMahon AP. Dorsalizing signal Wnt-7a required for normal polarity of D-V and A-P axes of mouse limb. Nature 1995;374:350–3.

Pende M, Vadiwala K, Schmidbaur H, Stockinger AW, Murawala P, Saghafi S, et al. A versatile depigmentation, clearing, and labeling method for exploring nervous system diversity. Sci Adv 2020;6:eaba0365.

Perez G, Barber GP, Benet-Pages A, Casper J, Clawson H, Diekhans M, et al. The UCSC Genome Browser database: 2025 update. Nucleic Acids Res 2025;53:D1243–9.

Pollard KS, Hubisz MJ, Rosenbloom KR, Siepel A. Detection of nonneutral substitution rates on mammalian phylogenies. Genome Res 2010;20:110–21.

Reifers F, Böhli H, Walsh EC, Crossley PH, Stainier DY, Brand M. Fgf8 is mutated in zebrafish acerebellar (ace) mutants and is required for maintenance of midbrain-hindbrain boundary development and somitogenesis. Development 1998;125:2381–95.

Riddle RD, Ensini M, Nelson C, Tsuchida T, Jesse ILl TM, Tabin” C. Induction of the LIM Homeobox Gene Lmx7 by WNT7a Establishes Dorsoventral Pattern in the Vertebrate Limb. Cell 1995;83:631–40.

Sandelin A, Alkema W, Engström P, Wasserman WW, Lenhard B. JASPAR: an openaccess database for eukaryotic transcription factor binding profiles. Nucleic Acids Res 2004;32:D91–4.

Schibler A, Malicki J. A screen for genetic defects of the zebrafish ear. Mech Dev 2007;124:592–604.

Schindelin J, Rueden CT, Hiner MC, Eliceiri KW. The ImageJ ecosystem: An open platform for biomedical image analysis. Mol Reprod Dev 2015;82:518–29.

Schweizer H, Johnson RL, Brand-Saberi B. Characterization of migration behavior of myogenic precursor cells in the limb bud with respect to Lmx1b expression. Anat Embryol 2004;208:7–18.

Seger C, Hargrave M, Wang X, Chai RJ, Elworthy S, Ingham PW. Analysis of Pax7 expressing myogenic cells in zebrafish muscle development, injury, and models of disease. Dev Dyn 2011;240:2440–51.

Sentmanat MF, Peters ST, Florian CP, Connelly JP, Pruett-Miller SM. A survey of validation strategies for CRISPR-Cas9 editing. Sci Rep 2018;8:888.

Siepel A, Bejerano G, Pedersen JS, Hinrichs AS, Hou M, Rosenbloom K, et al. Evolutionarily conserved elements in vertebrate, insect, worm, and yeast genomes. Genome Res 2005;15:1034–50.

Stebbins GL, Basile DV. Phyletic phenocopies: A useful technique for probing the genetic and developmental basis of evolutionary change. Evolution 1986;40:422–5.

Stewart TA, Lemberg JB, Taft NK, Yoo I, Daeschler EB, Shubin NH. Fin ray patterns at the fin-to-limb transition. Proc Natl Acad Sci U S A 2019. 10.1073/pnas.1915983117.

Stout CC, Tan M, Lemmon AR, Lemmon EM, Armbruster JW. Resolving Cypriniformes relationships using an anchored enrichment approach. BMC Evol Biol 2016;16:244.

Tahara Y. Normal stages of development in the lamprey, Lampetra reissued (dybowski). Zool Sci 1988;5:109–18.

Uemura O, Okada Y, Ando H, Guedj M, Higashijima S-I, Shimazaki T, et al. Comparative functional genomics revealed conservation and diversification of three enhancers of the isl1 gene for motor and sensory neuron-specific expression. Dev Biol 2005;278:587–606.

Vogel A, Rodriguez C, Warnken W, Izpisúa Belmonte JC. Dorsal cell fate specified by chick Lmx1 during vertebrate limb development. Nature 1995;378:716–20.

Wang Y, Shen Y, Feng C, Zhao K, Song Z, Zhang Y, et al. Mitogenomic perspectives on the origin of Tibetan loaches and their adaptation to high altitude. Sci Rep 2016;6:29690.

Wheeler TJ, Eddy SR. nhmmer: DNA homology search with profile HMMs. Bioinformatics 2013;29:2487–9.

Willis J, Burt de Perera T, Newport C, Poncelet G, Sturrock CJ, Thomas A. The structure and function of the sucker systems of hill stream loaches. bioRxiv 2019:851592. 10.1101/851592.

Yang Y, Niswander L. Interaction between the signaling molecules WNT7a and SHH during vertebrate limb development: dorsal signals regulate anteroposterior patterning. Cell 1995;80:939–47.

Yashima Y, Okada R, Kitagawa T. Differences in sexual morphological dimorphisms between two loach species of the genus Misgurnus (Cypriniformes: Cobitidae) in the River Shono system, Fukui Prefecture, Japan. J Vertebr Biol 2023;72:23035.1–14.

Zdral S, Bordignon SG, Meyer A, Ros MA, Woltering JM. Dorsoventral limb patterning in paired appendages emerged via regulatory repurposing of an ancestral posterior fin module. bioRxiv 2025:2025.04.16.648507. 10.1101/2025.04.16.648507.

